# Low-load blood flow restriction training and ischemia modulate expression of Na^+^,K^+^-ATPase and FXYDs in human skeletal muscle

**DOI:** 10.64898/2025.12.24.696414

**Authors:** Vid Jan, Katarina Miš, Alan Kacin, Anja Vidović, Tina Tomc Žargi, Klemen Stražar, Matej Podbregar, Tomaž Marš, Matej Drobnič, Alexander V. Chibalin, Sergej Pirkmajer

**Affiliations:** Institute of Pathophysiology, Faculty of Medicine, University of Ljubljana, Ljubljana, Slovenia; Department of Physiotherapy, Faculty of Health Sciences, University of Ljubljana, Ljubljana, Slovenia; Department of Orthopaedics, University Medical Centre Ljubljana, Ljubljana, Slovenia; General and Teaching Hospital Celje, Celje, Slovenia; Department of Orthopaedics, Faculty of Medicine, University of Ljubljana, Ljubljana, Slovenia; Integrative Physiology, Department of Molecular Medicine and Surgery, Karolinska Institutet, Stockholm, Sweden

**Keywords:** ACL injury rehabilitation, FXYD proteins, blood flow restriction (BFR), muscle deconditioning prevention, semitendinosus muscle, Na^+^, K^+^-ATPase

## Abstract

Anterior cruciate ligament (ACL) rupture leads to muscle deconditioning and downregulation of Na^+^,K^+^-ATPase (NKA). Low-load blood flow restriction (LL-BFR) training was shown to improve muscle function after ACL injury, but its effects on NKA are unknown. We analysed expression of NKA and its FXYD regulators in knee muscles from ACL-injured subjects undergoing LL-BFR, low-load training with sham blood flow restriction (LL-Sham), or no training (Control). Additionally, we dissected effects of ischemia components by subjecting cultured human myotubes to glucose deprivation and/or hypoxia. The LL-BFR group had higher vastus lateralis mRNA levels of NKAα1, NKAβ3, and FXYD5 than the LL-Sham group. In vitro, NKAα1, NKAβ1, and NKAβ3 mRNA and NKAα1 and NKAβ1 protein levels were downregulated by sustained ischemia and glucose deprivation, but not hypoxia, while FXYD5 protein was upregulated by glucose deprivation. Conversely, intermittent ischemia had no effect on NKA or FXYD expression. In conclusion, our study shows that LL-BFR training induces specific transcriptional adaptations in vastus lateralis after ACL injury, potentially contributing to functional improvements. Moreover, it shows that glucose availability plays a major role in modulating NKA and FXYD expression in muscle cells under ischemic conditions.

**NEW & NOTEWORTHY:**

- This is the first study exploring effects of low-load BFR training on Na^+^,K^+^-ATPase (NKA) and FXYD expression in knee muscles after anterior cruciate ligament injury.
- Effects of ischemia components were analysed by subjecting cultured human myotubes to glucose deprivation and/or hypoxia.
- LL-BFR training induces specific transcriptional adaptations in vastus lateralis after ACL injury, potentially contributing to functional improvements.
- Glucose plays a major role in modulating NKA and FXYD expression in cultured myotubes under ischemic conditions.

## INTRODUCTION

Downregulation of Na^+^,K^+^-ATPase (NKA) in skeletal muscle is a hallmark of physical inactivity (1) and was also observed in vastus lateralis muscle of subjects with anterior cruciate ligament (ACL) rupture (2). Although the knee can tolerate low mechanical loads after the ACL rupture, low-load resistance training alone is insufficient to prevent muscle deconditioning (3–5). However, evidence suggests that combining low-load resistance training with blood flow restriction (LL-BFR), achieved with inflatable tourniquet cuffs (4, 6–8), can counteract atrophy and loss of function of knee extensors following ACL rupture (5, 9–11). While BFR was shown to promote muscle NKA expression in healthy subjects (12, 13), it remains unknown whether similar effects occur after ACL injury.

Na^+^,K^+^-ATPase (NKA) transports two K^+^ ions into the cell and three Na^+^ ions out of the cell (14–16), thereby maintaining muscle excitability and contractility (17, 18). The core functional unit of NKA is a heterodimer that consists of the catalytic α-subunit (NKAα) and the non-catalytic β-subunit (NKAβ) (reviewed in (7, 19)). The NKAα subunit, a 100-112 kDa type IIC P-ATPase, has four isoforms (α1-4), of which three (α1-3) are expressed in skeletal muscle ((20), reviewed in (19, 21)). The NKAβ subunit, an essential 35-60 kDa glycoprotein (7, 22, 23), has three (β1-3) isoforms, all of which are expressed in skeletal muscle ((20), reviewed in (19, 21)). While NKAα/β-heterodimers are fully functional, their properties can be modulated by small transmembrane regulators of ion transport from the FXYD family (FXYD1-7)(24) (reviewed in (7, 8, 25)), among which FXYD1 (phospholemman) and FXYD5 (dysadherin) are most relevant in skeletal muscle (26–28).

Total NKA content (1, 29) determines skeletal muscle’s capacity to transport Na^+^ and K^+^, which in turn influences its resistance to fatigue (30). NKAα1, comprising ∼15-40% of muscle α-isoforms, is important for ion homeostasis in resting muscle and may also contribute to maintenance of muscle size (31). In contrast, NKAα2, comprising ∼60-85% of α-isoforms, plays a major role during muscle contractions ((32–35), reviewed in (21)). In healthy subjects, BFR combined with cycling was shown to increase the abundance of NKAα1, NKAβ1, and/or FXYD1 in a fibre type-dependent manner (12), while LL-BFR training using a knee extension apparatus increased the abundance of NKAα2 and NKAβ1 (13). Conversely, subjects with ACL rupture who did not undergo LL-BFR training had reduced total NKA content and lower NKAα2 abundance in vastus lateralis muscle (2). These findings led us to explore whether upregulation of NKA might have contributed to improved function of knee extensors observed in subjects with ACL rupture who performed LL-BFR training in our previous study (5).

BFR restricts the delivery of oxygen and nutrients, such as glucose, to working skeletal muscle, resulting in energy stress (36–40). Under these ischemic conditions, activation of AMP-activated protein kinase (AMPK), a cellular energy sensor, likely contributes to adaptive responses in skeletal muscle (41–43). AMPK may regulate NKA activity and content through several mechanisms (reviewed in (21, 44)), including by promoting dephosphorylation of NKAα1 at Ser^18^ (45) and/or at Tyr^10^ (46), as well as by increasing phosphorylation of FXYD1 at Ser^68^ (47). Our recent work demonstrated that both AMPK activation and glucose deprivation both modulated expression of NKA subunits and FXYDs in cultured myotubes under normoxic conditions (48). However, it remains unclear whether and how glucose deprivation and hypoxia, which occur together during ischemia, interact to regulate NKA or FXYDs.

In the present study, we used samples from our previous work (5) to assess whether LL-BFR training altered expression of NKA or FXYDs in vastus lateralis muscle of subjects with ACL rupture. Additionally, to separately evaluate response of NKA and FXYDs to hypoxia and/or glucose deprivation, the key components of ischemia, we used cultured human myotubes.

## MATERIALS AND METHODS

### Ethical authorization

The Republic of Slovenia National Medical Ethics Committee approved the training study in subjects with an ACL injury (KME 45/08/14, amendment no.: 62/05/12) (5) and the use of primary human skeletal muscle cells (HSMC), obtained from a separate group of subjects with an ACL injury (ethical approvals no.: 71/05/12 and 0120-698/2017/4). Participation in the study was voluntary, and all subjects had given written informed consent prior to enrolment.

### Materials

Cell culture flasks, plates and Petri dishes were from Sarstedt or TPP. Advanced MEM, DMEM, foetal bovine serum (FBS), MEM Vitamin solution (100X), GlutaMAX (100X), Fungizone (250 µg/mL of amphotericin B), gentamicin (10 mg/mL), Pierce BCA Protein Assay Kit, Pierce Enhanced Chemiluminescence (ECL) Western Blotting Substrate, High-Capacity cDNA Reverse Transcription Kit, MicroAmp optical 96-well reaction plates, MicroAmp optical adhesive foils and TaqMan Universal Master Mix were from Thermo Fisher Scientific. 4–12% Criterion XT Bis-Tris polyacrylamide gels, XT MES electrophoresis buffer, goat anti-mouse IgG – horseradish peroxidase conjugate, and goat anti-rabbit IgG – horseradish peroxidase conjugate were from Bio-Rad. Amersham ECL Full-Range Rainbow Molecular Weight Markers were from GE Healthcare Life Sciences. The polyvinylidene fluoride (PVDF) membrane was from Merck Millipore. CP-BU NEW X-ray films were from AGFA HealthCare. RNeasy Plus Mini Kit and RNeasy Fibrous Tissue Mini Kit were from Qiagen. All other reagents, unless otherwise stated, were from Sigma-Aldrich and of analytical grade.

### Test subjects and training programme

Samples of vastus lateralis and semitendinosus muscles, which we analysed in the current study, were obtained during surgery from subjects with ACL rupture as part of the previously published study (5). The current study presents only new data, with no overlap with results previously published (5). Characteristics of the subjects and the study protocol were reported in detail (5). Briefly, the study was designed as a prospective, single-centre, single-blinded, quasi-randomised trial controlled with a sham intervention. Recruitment took place between May 2015 and December 2016 at the Department of Orthopaedics, University Medical Centre Ljubljana. Subjects were allocated to the groups in a concealed manner by contacting the holder of the allocation plan, who was not on site. Inclusion criteria were age of subjects between 18 and 45 years, complete unilateral ACL rupture diagnosed by MRI and clinical assessment, no previous injury or surgery of the affected knee, no neurologic deficits on the lower limbs, no systemic diseases, no previous injuries to the contralateral knee, pain intensity during exercise of less or equal to 2 according to the visual analogue scale, sufficient range of motion of the affected knee for the subjects to perform exercise (active extension deficit ≤ 5°, active flexion ≥ 120°).

Exclusion criteria were concomitant intra-articular pathology that would compromise the safety or tolerability of exercise (i.e. significant damage to articular cartilage or menisci), spine or other lower limb injuries, neuromuscular impairment, presence or history of vascular disease or deep vein thrombosis. Of the 35 subjects screened for eligibility, 16 met the criteria and 12 were included in the study. The 12 subjects (6 women and 6 men) completed the entire protocol and were included in the data analysis. The gender distribution into low-load training with sham blood flow restriction (LL-Sham) or low-load training with true blood flow restriction (LL-BFR) was even (i.e. 3 males and 3 females in each group). Subjects were blinded to the actual physiological effects of the different pneumatic cuff pressures used during the training intervention. A group (n = 6) of subjects with an ACL rupture and equal physical characteristics who did not perform any training and were not subjected to blood flow restriction was included as an additional control.

The 3-week LL-BFR training protocol for the injured leg was designed to induce skeletal muscle hypertrophy and improve muscle performance (36, 49). The training intervention comprised 9 exercise sessions (Fig. 1A), which were performed with either LL-Sham or LL-BFR in the thigh muscles. Every session included 4 sets of knee flexion and extension exercises performed at 40-repetition maximum (40RM) to volitional failure with the injured leg only. A 13.5 cm wide pneumatic double-chamber cuff with asymmetric pressure (Ischemic Trainer, University of Ljubljana and Iskra Medical d.o.o.) was applied to the proximal thigh to induce a partial blood flow restriction in the thigh muscles. After the initial warm-up consisting of 10–12 repetitions at minimal workload, the cuff was inflated with air to a cumulative pressure of 150 mmHg (inner cuff compartment = 185 mmHg; outer cuff compartment = 125 mmHg) (6) and left on a resting muscle for 30 seconds.

**Fig. 1.**
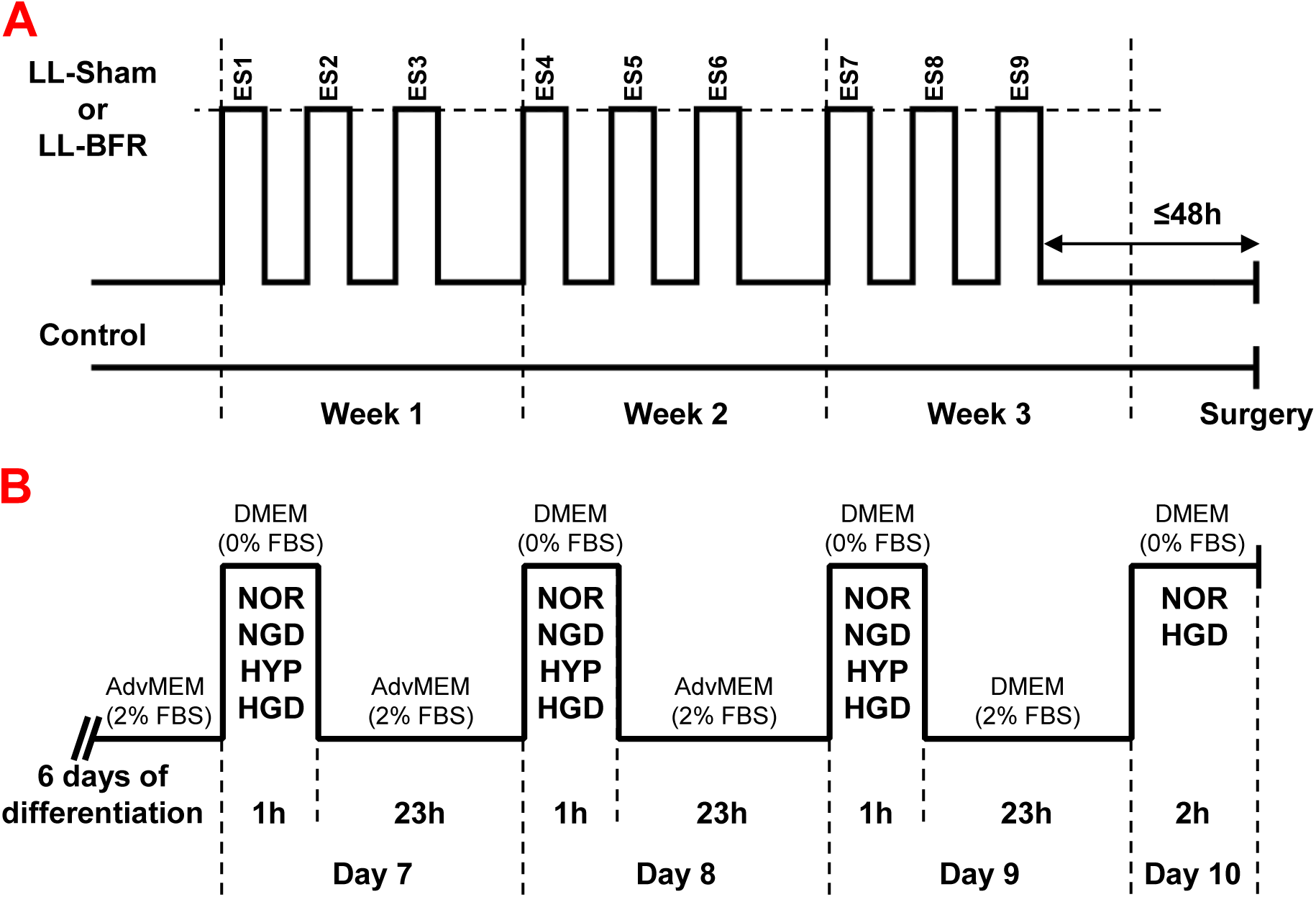
Schematic representation of the training protocol and *in vitro* preconditioning with intermittent exposure to ischemia, hypoxia, or glucose deprivation. (A) Training intervention comprised 9 exercise sessions (ES) that were performed with either true blood flow restriction (LL-BFR) or sham blood flow restriction (LL-Sham). Subjects in the control group did not perform any interventions (Control). Every session included 4 sets of knee flexion and extension exercises performed at 40RM to volitional failure with injured leg only. Muscle samples were procured at the time of hamstring tendon harvesting, which is performed in the early stages of surgery (max 20 min after incision). Once collected, muscle samples were immediately frozen in liquid nitrogen. See text for further details. (B) Myotubes were differentiated in Advanced MEM (AdvMEM) with 2% FBS for 6 days. On days 7-9, they were exposed to normoxia (NOR), glucose deprivation (glucose-free medium, NGD), hypoxia (0.1% O_2_, HYP), or both – ischemia (HGD) in serum-free DMEM for 1 hour/day. After the last preconditioning myotubes were switched to DMEM with 2% FBS for 23 hours and then exposed to ischemia (HGD) or NOR for 2 hours. See text for details.

Participants in the LL-BFR group consistently completed their training sessions prior to their matched peers in the LL-Sham group. Each LL-BFR training session involved four sets of unilateral knee extensions and flexions under ischemic conditions, with a cuff pressure of 150 mmHg. These exercises were performed with the injured leg only, at the participant’s initial 40RM, each set continuing to volitional failure. The workload was held constant throughout the intervention period.

The LL-Sham group executed an identical unilateral exercise protocol with the injured leg, with the key difference being that the cuff was inflated to only 20 mmHg. To ensure volume equivalence between groups, each participant in the sham group replicated the exact number of repetitions performed in each set by their paired counterpart in the LL-BFR group during the corresponding session.

Participants were paired across groups based on gender, Lysholm score, and body mass index (BMI). The volume of each training session was calculated by summing the total work (repetitions × load) for both the extensor and flexor muscle groups. The cumulative training dose was computed as the sum of volumes across all sessions. This approach ensured nearly identical mechanical workload and hence mechanical muscle strain in both groups.

The subjects were assigned to the LL-BFR and LL-Sham groups in a counterbalanced manner. The first two subjects were assigned to the LL-BFR group and the next two to the LL-Sham group, which was then repeated until both groups were complete. The matching of the subjects across the groups according to gender, BMI, and Lysholm score was performed by a physiotherapist not directly involved in the study. This ensured that the most important baseline measures and prognostic indicators ware consistent between the groups. Subjects were asked not to change their regular daily activity routine during the intervention and to keep a daily diary of all their physical activities. Subjects were familiarised with the testing and training protocols during preliminary visits to reduce the impact of the initial motor learning on the test results. The interval was standardized, all subjects were familiarized with the protocols 48 hours prior to the initial measurements. Particular emphasis was placed on acquiring the correct technique and pace for weightlifting. They were also familiarised with the use of a pneumatic tourniquet during exercise. All subjects received the treatment or control conditions as assigned.

### Skeletal muscle sampling

Skeletal muscle samples in the LL-BFR and LL-Sham group were collected during surgery from the trained leg within 48 hours after the last exercise session (Fig. 1A). The samples were collected at the beginning of the surgical ACL reconstructions (maximally 20 minutes after incision without tourniquet closure), which were performed under spinal block anaesthesia. One sample was surgically cut from semitendinosus muscle during harvesting the autologous graft for ACL reconstruction, while the other was taken from the lateral part of vastus lateralis muscle with a percutaneous needle biopsy technique using a 50 mm Bergström needle with syringe suction. Once collected, muscle samples were immediately frozen in liquid nitrogen in the operating theatre at the Department of Orthopaedics. After freezing, samples were transferred in liquid nitrogen to the Institute of Pathophysiology, where they were stored at -80°C until further processing.

### Cultures of primary human skeletal muscle cells

Human skeletal muscle cells (HSMC) were obtained from samples of semitendinosus muscle that are routinely removed and discarded as surgical waste during ACL reconstruction, as described (48, 50–53). These samples were collected from a separate group of subjects who had not undergone training prior to reconstructive surgery and were not included in the training study described above. Briefly, once surgically cut from semitendinosus muscle during harvesting the autologous graft for ACL reconstruction, muscle samples were immediately stored in sterile growth medium (Advanced MEM supplemented with 10% (v/v) FBS, 1% (v/v) 100X MEM vitamin solution, 1% (v/v) 100X GlutaMAX, 0.3% (v/v) Fungizone (0.75 μg/mL of amphotericin B) and 0.15% (v/v) gentamicin (15 μg/mL)) and transferred to the skeletal muscle cell laboratory at the Institute of Pathophysiology. The muscle tissue was cleaned of connective and adipose tissue, cut into small pieces and trypsinized. Primary cultures were grown at 37°C in a humidified atmosphere (5% CO_2_) in the growth medium. Before reaching confluence, skeletal muscle cells were isolated using MACS CD56 MicroBeads (Miltenyi Biotec), as described in detail (53). Purified HSMC were cultured in growth medium for 2–3 passages before being used for the experiments.

### Sustained (24-hour) exposure to ischemia, hypoxia, and/or glucose deprivation

HSMC were differentiated into myotubes by reducing the FBS concentration from 10% (v/v) to 2% (v/v) for 7–9 days, as described(53). One day before the experiment, myotubes were switched to DMEM (1 g/L glucose) with 2% (v/v) FBS for 24 hours. On the last day, myotubes were exposed to control conditions (serum-free DMEM with 1 g/L glucose, NOR), glucose deprivation (serum-free DMEM without glucose, NGD), hypoxia (0.1% O_2_, serum-free DMEM with 1 g/L glucose, HYP), or both – ischemia (0.1% O_2_ and serum-free DMEM without glucose, HGD). Hypoxic conditions (0.1% O_2_) were established in a New Brunswick Galaxy 48R CO_2_ incubator (Eppendorf, Germany).

### Preconditioning with intermittent exposure to ischemia, hypoxia, and/or glucose deprivation

HSMC were differentiated into myotubes for 6 days in Advanced MEM with 2% FBS (Fig. 1B), as described (53). On days 7, 8, and 9, HSMC were subjected to preconditioning with glucose deprivation, hypoxia, or both for 1 hour/day. Before treatment, myotubes were washed with glucose- and serum-free DMEM and then placed in serum-free DMEM with 1 g/L glucose in normoxia (NOR), serum-free DMEM without glucose in normoxia (NGD), serum-free DMEM with 1 g/L glucose in hypoxia (0.1% O_2_) (HYP), or serum-free DMEM without glucose in hypoxia (0.1% O_2_) (HGD, artificial ischemia) (Fig. 1B). Each treatment lasted 1 hour from the time the 0.1% O_2_ concentration was reached in the hypoxic incubator. On days 7 and 8, myotubes were placed back into Advanced MEM with 2% FBS. On day 9, myotubes were switched to DMEM with 1 g/L glucose and 2 % FBS for 23 hours (Fig. 1B), after which they were exposed to artificial ischemia (HGD, serum-free DMEM without glucose, 0.1% O_2_) or normoxic control conditions (NOR, serum-free DMEM with 1g/L glucose) for 2 hours.

### Immunoblotting

The frozen muscle samples were first ground to a fine powder in liquid nitrogen using a pestle and mortar. They were then homogenised with a mechanical homogeniser and the homogenisation buffer (137 mM NaCl, 2.7 mM KCl, 1 mM MgCl_2_, 1% (v/v) Triton X-100, 10% (w/v) glycerol, 20 mM Tris (pH 7.8), 10 mM NaF, 1 mM EDTA, 0.5 mM Na_3_VO_4_, 1 mM phenylmethylsulfonyl fluoride, 1% (v/v) protease inhibitor cocktail). In the next step, the samples were put on a turning wheel for 1 hour at 4°C. At the end of the *in vitro* experiments, the myotubes were washed with ice-cold phosphate-buffered saline (PBS: 137 mM NaCl, 2.7 mM KCl, 10 mM Na_2_HPO_4_, 1.8 mM KH_2_PO_4_, pH 7.4) and lysed in the homogenisation (the prolonged ischemia experiment) or Laemmli buffer (the intermittent ischemia experiment).

The muscle and myotube lysates prepared in the homogenization buffer were centrifuged at 12000×g for 15 min. Protein concentrations in the supernatant were measured using the BCA protein assay. Samples of vastus lateralis and semitendinosus muscle were equalized to the same final protein concentrations (separately for each muscle because the size of samples of vastus lateralis muscle was much smaller than those of semitendinosus) and mixed with Laemmli buffer (62.5 mM Tris-HCl (pH 6.8), 2% (w/v) sodium dodecyl sulfate (SDS), 10% (w/v) glycerol, 5% (v/v) 2-mercaptoethanol, 0.002% (w/v) bromophenol blue). Equal amount of total protein was loaded for lysates prepared from vastus lateralis (5 μg/well) and semitendinosus (35 or 70 μg/well) muscles. Due to different loading, immunoblotting analyses of the vastus lateralis and semitendinosus samples were separate. Pierce 660-nm Protein Assay, which is compatible with the Laemmli buffer, was used to measure total protein concentrations in myotube lysates that were prepared directly in the Laemmli buffer. Total protein concentrations of myotube lysates were equalized for each donor separately, but equal amount of total protein was loaded for each donor (range: 12–35 μg/well).

Proteins were resolved by SDS-PAGE (4–12% polyacrylamide gels) at 200 V for 40 minutes using the Criterion system and XT MES buffer. Proteins were transferred to PVDF membrane with wet electrotransfer at 100 V (transfer buffer: 31 mM Tris, 0.24 M glycine, 10% methanol and 0.01% SDS). Ponceau S staining (0.1% in 5% acetic acid) staining was used as an additional control of the uniformity of sample loading and quality of transfer. The membranes were then blocked with 7.5% dry skimmed milk in Tris-buffered saline with Tween (TBST: 20 mM Tris, 150 mM NaCl, 0.02% Tween 20, pH 7.5) for 1 hour at room temperature. The blocked membranes were incubated with primary antibodies (in 20 mM Tris, 150 mM NaCl, pH 7.5, 0.1% BSA, and 0.1% sodium azide) overnight at 4°C. After washing with TBST, the membranes were incubated with secondary antibody-horseradish peroxidase conjugate in TBST containing 5% (w/v) dry skimmed milk for 1 hour at room temperature. Enhanced chemiluminescence was used to visualise the immunoreactive bands on X-ray films. Quantification was performed using the GS-800 Densitometer (Bio-Rad) and Quantity One 1-D Analysis Software (Bio-Rad).

Primary antibodies that were used in this study are presented in Table 1. Briefly, the monoclonal antibodies against NKAβ1 (MA3-930) (48) were from Thermo Fisher Scientific. Monoclonal antibodies against NKAα1 (05–369) (46, 48, 53) and polyclonal antibodies against NKAα2 (AB9094)(48, 53) were from Merck Millipore. Polyclonal antibodies against AMPKα (cs-2532)(48), monoclonal antibodies against phospho-AMPKα (Thr^172^) (cs-2535) (46, 48, 54), monoclonal antibodies against acetyl-CoA carboxylase (cs-3676) (46, 48), polyclonal antibodies against phospho-acetyl-CoA carboxylase (Ser^79^) (cs-3661) (48, 54), monoclonal antibodies against phospho-NKAα1 (Tyr^10^) (E1Y9C) (cs-13566) (46) and monoclonal antibodies against Sp1 (D4C3) (cs-9389) (53) were from Cell Signaling Technology. Polyclonal antibodies against phospho-FXYD1 (Ser^68^) (PAB0389) (53, 55) were from Abnova, polyclonal antibodies against FXYD1 (13721-1-AP) (48, 53) were from Proteintech, polyclonal antibodies against hypoxia-inducible factor-1α (HIF-1α) (50, 54, 56, 57) were from Novus Biologicals, while polyclonal antibodies for FXYD5 were a kind gift from Dr. Haim Garty (Weizmann Institute of Science, Rehovot, Israel) (28, 53, 58).

**Table 1.** Antibodies used for immunoblotting

| Target | Primary antibody |  |  | Secondary antibody |  |
| --- | --- | --- | --- | --- | --- |
|  | Species (clonality) | Dilution | Cat. No. | Species | Dilution |
| ACC | rabbit (M) | 1:1000 | #3676 (CST) | GAR | 1:25000 |
| AMPKα | rabbit (P) | 1:1000 | #2532 (CST) | GAR | 1:10000 |
| FXYD1 | rabbit (P) | 1:500 | #13721-1-AP (PT) | GAR | 1:10000 |
| FXYD5 | mouse (P) | 1:500 | See text. | GAM | 1:25000 |
| HIF-1α | rabbit (P) | 1:1000 | NB100-449 (NB) | GAR | 1:6250 |
| NKAα1 | mouse (M) | 1:2000 | #05-369 (Merck) | GAM | 1:25000 |
| NKAα2 | rabbit (P) | 1:4000 | #AB9094 (Merck) | GAR | 1:50000 |
| NKAβ1 | mouse (M) | 1:2000 | MA3-930 (TFS) | GAM | 1:100000 |
| pACC Ser<sup>79</sup> | rabbit (P) | 1:1000 | #3661 (CST) | GAR | 1:10000 |
| pAMPKα Thr<sup>172</sup> | rabbit (M) | 1:1000 | #2535 (CST) | GAR | 1:25000 |
| pFXYD1 Ser<sup>68</sup> | rabbit (P) | 1:500 | PAB0389 (AB) | GAR | 1:10000 |
| pNKAα1 Tyr<sup>10</sup> | rabbit (M) | 1:1000 | #13566 (CST) | GAR | 1:10000 |
| SP1 | rabbit (M) | 1:500 | #9389 (CST) | GAR | 1:20000 |

*Abbreviations:* M – monoclonal; P – polyclonal; CST – Cell Signaling Technology; PT – Proteintech; NB – Novus Biologicals; TFS – Thermo Fisher Scientific; AB – Abnova; GAR – goat anti-rabbit IgG-HRP conjugate #1706515 from Bio-Rad; GAM – goat anti-mouse IgG- HRP conjugate #1706516 from Bio-Rad.

Three out of the six lysates prepared from semitendinosus muscle samples for immunoblotting contained low levels of myosin heavy chains, indicating they may have consisted mainly of fibrous tissue rather than skeletal muscle fibres. This was evident on Ponceau S-stained membranes (see Raw data of Immunoblotting) and during immunoblotting with antibodies specific for myosin heavy chains (data not shown). Although the results from these samples did not qualitatively differ (data not shown), only the three samples of semitendinosus muscle with prominent myosin heavy chain expression were included in the final immunoblotting analysis. Due to small size of vastus lateralis biopsies, we were able to perform analyses on protein samples from of 5 donors from the LL-BFR group. Due to low amount of protein that we were able to load, some of the target proteins in lysates from vastus lateralis muscles were below detection level, which is why *n* is less than 5 for some of the densitometric analyses.

### Quantitative real-time polymerase chain reaction

Total RNA from muscle tissue was extracted using the RNeasy Fibrous Tissue Mini Kit, while RNA from cultured myotubes was extracted using the RNeasy Plus Mini Kit. Total RNA was reverse transcribed into cDNA using the High-Capacity cDNA Reverse Transcription Kit. Quantitative real-time polymerase chain reaction (PCR) was performed with the 7500 Real-Time PCR System (Applied Biosystems, Thermo Fisher Scientific) using the TaqMan Universal Master Mix and TaqMan gene expression assays: ATP1A1 (Hs00167556_m1, ATP1A2 (Hs00265131_m1), ATP1A3 (Hs00958036_m1), ATP1B1 (Hs00426868_g1), ATP1B2 (Hs00155922_m1), ATP1B3 (Hs00740857_mH), FXYD1 (Hs00245327_m1), FXYD5 (Hs00204319_m1), VEGFA (Hs00900055_m1), PGK1 (Hs00943178_g1), HIF1A (Hs00153153_m1), and 18S rRNA (Hs99999901_m1). The mRNA expression levels are reported as gene expression ratios (target gene mRNA/18S rRNA) according to the following equation: (1+ E)^Ct,18S^ ^rRNA^/(1 + E)^Ct,target^ ^mRNA^, where E is the efficiency of PCR, while C_t_ is the threshold cycle for the endogenous control (18S rRNA) or target genes(59, 60).

### Statistical analysis

The data are reported as mean values with standard deviation (SD). The statistical analysis of the differences between the groups was performed with GraphPad Prism 6 (GraphPad Software). Either the non-parametric Kruskal-Wallis test with Dunn’s post-hoc test for clinical samples or a one- or two-way ANOVA with Dunnett’s *post hoc* test for cell culture experiments was used. The difference was considered statistically significant if *P*<0.05.

## RESULTS

### Effect of LL-BFR and LL-Sham training on the expression of the NKA subunits, FXYD1, and FXYD5 in vastus lateralis and semitendinosus muscles

NKAα1 mRNA levels in vastus lateralis muscle were higher in the LL-BFR group than in the Control group (*P*=0.03) and LL-Sham group (*P*=0.04) (Fig. 2A), while protein levels were not statistically different (Fig. 2I; *P*=0.33, *P*>0.99, respectively). NKAα1 protein levels in semitendinosus muscle, but not mRNA levels (*P*=0.10), tended to be higher in the LL-BFR group than in the Control group (*P*=0.05) (Fig. 2I). NKAα2 expression in both muscles was similar in all three groups (Fig. 2B,J; all *P*≥0.14). NKAα3 mRNA levels in vastus lateralis muscle were not significantly different (all *P*≥0.08), but they were lower in semitendinosus muscle in the LL-Sham (*P*=0.006) and LL-BFR (*P*=0.03) groups (Fig. 2C). However, it is important to note that NKAα3 mRNA expression was very low and NKAα3 protein could not be detected reliably.

**Fig. 2.**
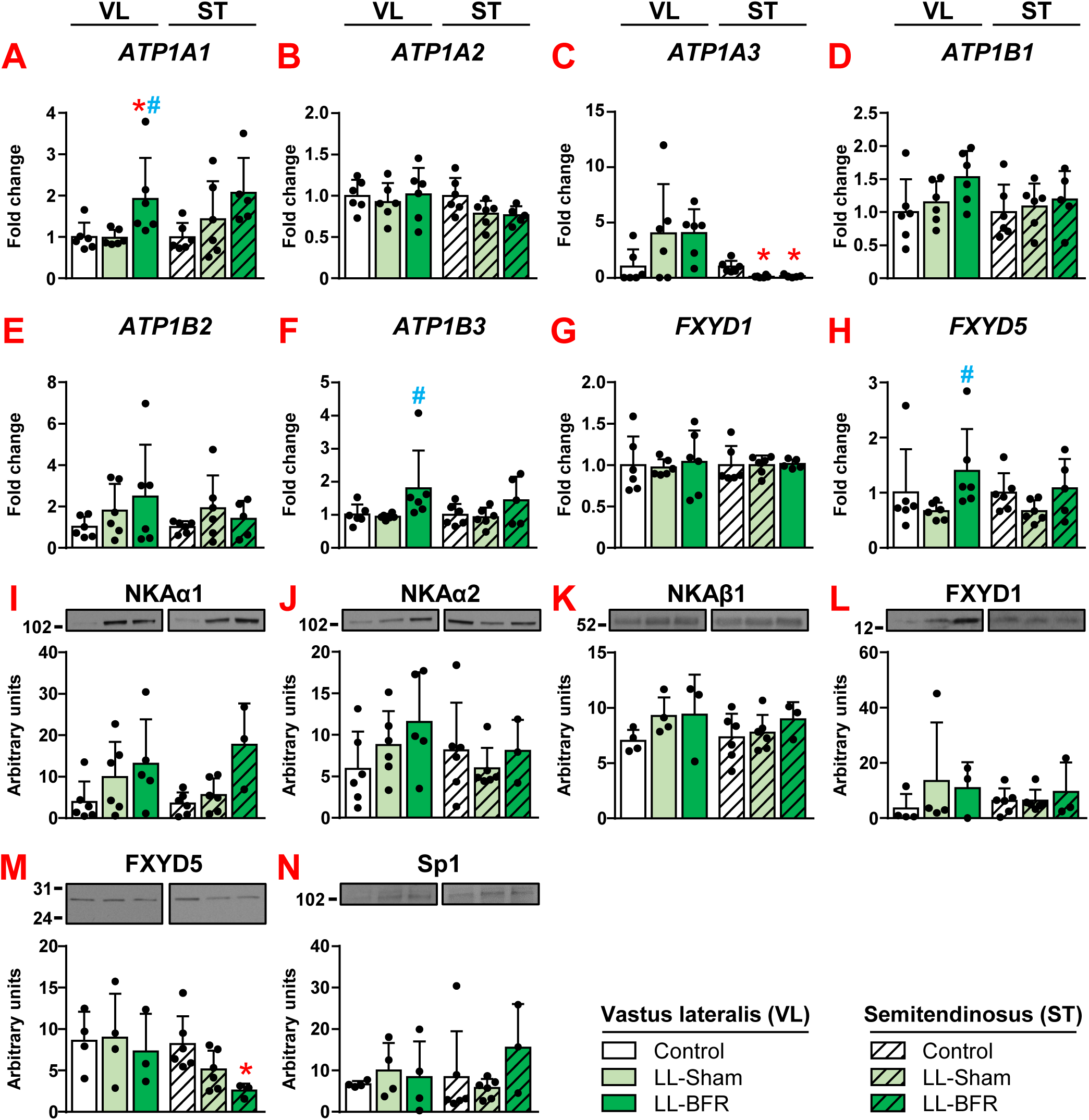
Effect of LL-BFR and LL-Sham training on the expression of NKA subunits, FXYD1, FXYD5 and Sp1 in vastus lateralis and semitendinosus muscles. qPCR was used to determine gene expression of (A) NKAα1 (*ATP1A1*), (B) NKAα2 (*ATP1A2*), (C) NKAα3 (*ATP1A3*), (D) NKAβ1 (*ATP1B1*), (E) NKAβ2 (*ATP1B2*), (F) NKAβ3 (*ATP1B3*), (G) *FXYD1*, and (H) *FXYD5*. 18S rRNA was used as endogenous control. Immunoblotting was used to estimate protein abundance of (I) NKAα1, (J) NKAα2, (K) NKAβ1, (L) FXYD1, (M) FXYD5, and (N) Sp1. Results are means ± SD (n=3-6); \**P*<0.05 vs. Control; #*P*<0.05 vs LL-Sham. Kruskal-Wallis with Dunn‘s post-hoc test was used for statistical evaluation.

In vastus lateralis muscle, NKAβ1 expression was similar among all groups (Fig. 2D,K; all *P*≥0.15), whereas NKAβ3 mRNA levels were higher in the LL-BFR group than in the LL-Sham group (*P*=0.03) (Fig. 2F). In semitendinosus muscle, NKAβ mRNA and protein levels were similar in all groups (Fig. 2D-F,K; all *P*≥0.63).

FXYD1 mRNA and protein levels were similar between groups in both vastus lateralis and semitendinosus muscles (Fig. 2G,L; all *P*≥0.60). In vastus lateralis muscle, FXYD5 mRNA expression was higher in the LL-BFR group than in the LL-Sham group (Fig. 2H; *P*=0.02), but protein levels did not differ (Fig. 2M; all *P*≥0.99). In semitendinosus muscle, FXYD5 protein levels were similar between the LL-BFR and LL-Sham group (*P*=0.74) but were lower in the LL-BFR than in LL-Control group (*P*=0.03) (Fig. 2M), with no difference in mRNA expression (Fig. 2H; all *P*≥0.18). The abundance of the transcription factor Sp1, which is involved in the regulation of NKA subunit expression (61) and responds to hypoxia (62), was similar across all groups (Fig. 2N; all *P*≥0.46).

### Effects of LL-BFR and LL-Sham training on AMPK signalling and the phosphorylation of NKAα1 (Tyr^10^) and FXYD1(Ser^68^) in vastus lateralis and semitendinosus muscles

AMPK activity, as indicated by phosphorylation of the catalytic AMPKα-subunit at Thr^172^ (Fig. 3A-C) and its substrate acetyl-CoA carboxylase (ACC) at Ser^79^ (Fig. 3D-F), was similar among the three groups (all *P*≥0.26). In semitendinosus muscle, phosphorylation of NKAα1 at Tyr^10^ tended to be lowest in the LL-BFR group compared to LL-Control group, but this difference was not statistically significant (Fig. 3G; *P*=0.17; Fig 3H; *P*=0.12). Phosphorylation of FXYD1 at Ser^68^ was also similar across all groups (Fig. 3I,J; all *P*≥0.92).

**Fig. 3.**
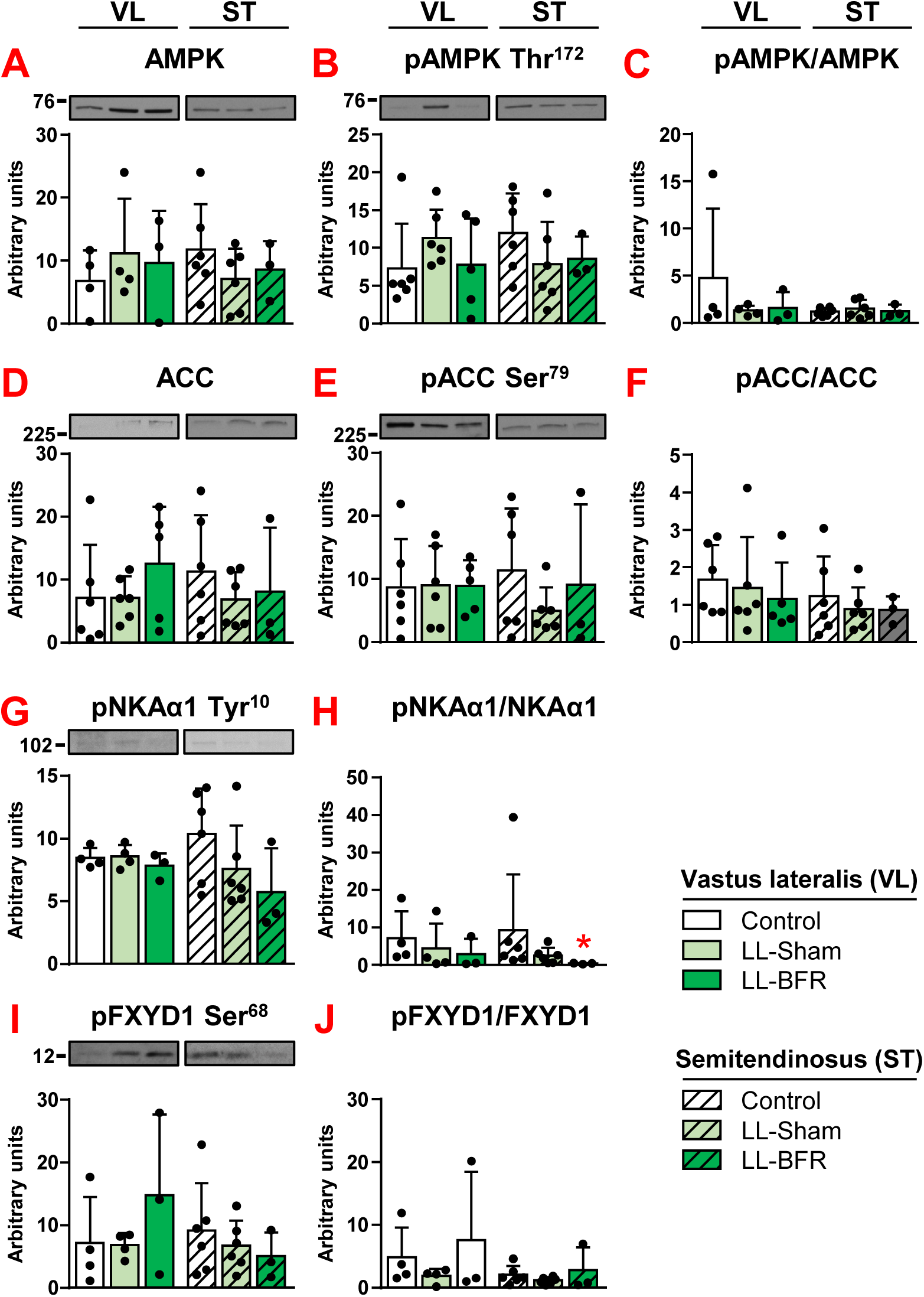
Effects of LL-BFR and LL-Sham training on AMPK signalling and the phosphorylation of NKAα1 (Tyr^10^) and FXYD1(Ser^68^) in vastus lateralis and semitendinosus muscles. Immunoblotting was used to estimate protein abundance of (A) AMPK, (B) pAMPK (Thr^172^), (C) phosphorylation of AMPK, relative to total protein (D) ACC, (E) pACC (Ser^79^), (F) phosphorylation of ACC, relative to total protein, (G) pNKAα1 (Tyr^10^), (H) phosphorylation of NKA α1, relative to total protein, (I) pFXYD1 (Ser^68^), and (J) phosphorylation of FXYD1, relative to total protein. Results are means ± SD (n=3-6); \**P*<0.05 vs. Control; #*P*<0.05 vs LL-Sham. Kruskal-Wallis with Dunn‘s post-hoc test was used for statistical evaluation.

### Effect of sustained (24-hour) ischemia on HIF-1α, AMPK, and NKA in cultured human myotubes

To separately dissect the role of hypoxia and glucose deprivation, the key components of ischemia, on NKA subunits and FXYD expression, cultured human myotubes were exposed to artificial (*in vitro*) ischemia (i.e. hypoxia (0.1% O_2_) and glucose deprivation), hypoxia (0.1% O_2_), or glucose deprivation alone (Fig. 4).

**Fig. 4.**
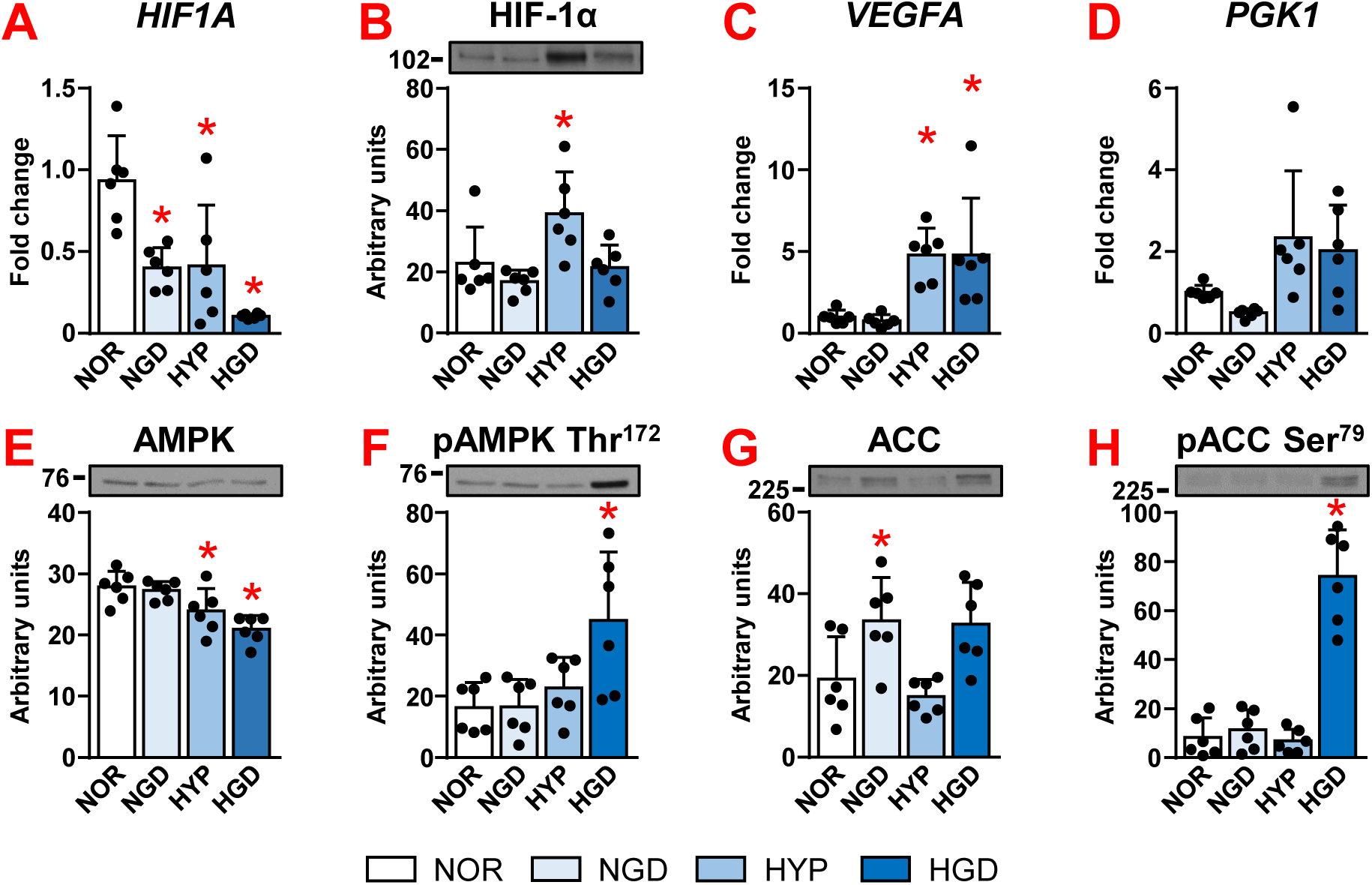
Effect of sustained (24-hour) ischemia on HIF-1α and AMPK in cultured human myotubes. For the last 24 hours differentiated human myotubes were exposed to normoxia (NOR), glucose deficiency (glucose-free medium, NGD), hypoxia (0.1% O_2_, HYP), or both – ischemia (HGD) in serum-free DMEM. See text for details. qPCR was used to estimate gene expression of (A) HIF-1α (*HIF1A*), (C) *VEGFA*, and (D) *PGK1*. 18S rRNA was used as endogenous control. Immunoblotting was used to estimate protein abundance of (B) HIF-1α, (E) AMPK, (F) pAMPK (Thr^172^), (G) ACC, and (H) pACC (Ser^79^). Results are mean ± SD for n=6; \**P*<0.05 vs. NOR. Ordinary one-way ANOVA with Dunnett’s post-hoc test was used for statistical evaluation.

The response to hypoxia was assessed by evaluating the expression of hypoxia-inducible factor-1α (HIF-1α), a key regulator of oxygen homeostasis (63), and its target genes phosphoglycerate kinase 1 (PGK1) and vascular endothelial growth factor A (VEGFA), as described (50). HIF-1α mRNA expression was suppressed by both hypoxia (*P*=0.003) and glucose deprivation (*P*=0.003) and reached the lowest level under ischemic conditions (*P*<0.001) (Fig. 4A). HIF-1α protein was upregulated in hypoxia (*P*=0.03) (Fig. 4B), which was accompanied by upregulation of VEGFA mRNA (*P*=0.007) (Fig. 4C). Ischemia enhanced down-regulation of HIF-1α mRNA and thereby probably blunted up-regulation of HIF-1α protein (Fig. 4B; *P*=0.99). A marked increase in the phosphorylation of AMPK (*P*=0.006) and ACC (*P*<0.001) (Fig. 4F,H) suggested ischemic myotubes were in energy stress.

Hypoxia alone did not significantly alter the expression of the NKA subunits, FXYD1, or FXYD5 (Fig. 5; all *P*≥0.15). Ischemia or glucose deprivation reduced the expression of NKAα1 (Fig. 5A,I; all *P*<0.001). Phosphorylation of NKAα1 at Tyr^10^ was reduced under ischemic conditions (*P*<0.001) (Fig. 5J). The mRNA expression of NKAα2 and NKAα3 remained unchanged under ischemic and glucose-deprived conditions (Fig. 5B,C,L; all *P*≥0.45). The NKAα3 protein is not shown because it could not be reliably detected in myotubes, most likely due to its very low expression. The transcription factor Sp1 was reduced under all three conditions and reached the lowest levels in ischemia (*P*≤0.004) (Fig. 5N).

**Fig. 5.**
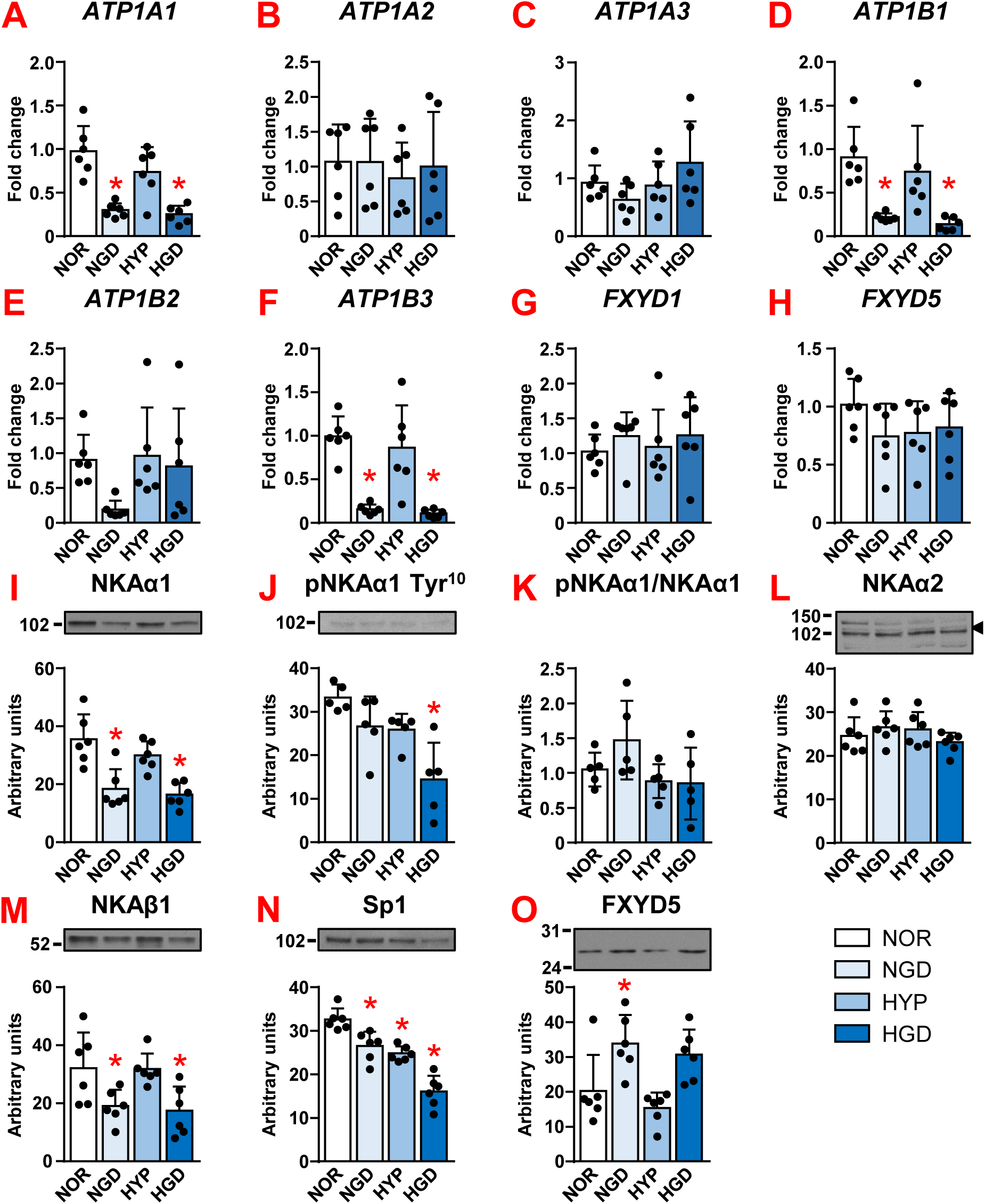
Effect of sustained (24-hour) ischemia on the expression of NKA subunits, FXYD1, and FXYD5 in cultured human myotubes. For the last 24 hours differentiated human myotubes were exposed to normoxia (NOR), glucose deficiency (glucose-free medium, NGD), hypoxia (0.1% O_2_, HYP), or both – ischemia (HGD) in serum-free DMEM. See text for details. qPCR was used to estimate gene expression of (A) NKAα1 (*ATP1A1*), (B) NKAα2 (*ATP1A2*), (C) NKAα3 (*ATP1A3*), (D) NKAβ1 (*ATP1B1*), (E) NKAβ2 (*ATP1B2*), (F) NKAβ3 (*ATP1B3*), (G) *FXYD1*, and (H) *FXYD5*. Immunoblotting was used to estimate protein abundance of (I) NKAα1, (J) pNKAα1 (Tyr^10^), (K) phosphorylation of NKAα1, relative to total protein, (L) NKAα2, (M) NKAβ1, (N) Sp1, and (O) FXYD5. Results are means ± SD for n=6 for qPCR and n=5-6 for immunoblotting; \**P*<0.05 vs. NOR. In case of unspecific bands arrowheads mark the specific bands that were used in densitometric analysis. Ordinary one-way ANOVA with Dunnett’s post-hoc test was used for statistical evaluation.

Ischemia and glucose-deprivation suppressed NKAβ1 mRNA and protein levels (*P*≤0.04) (Fig. 5D,M). While NKAβ2 mRNA expression was unaltered (Fig. 5E; all *P*≥0.11), NKAβ3 mRNA expression was almost completely suppressed by ischemia (*P*<0.001) or the lack of glucose (*P*<0.001) (Fig. 5F). FXYD1 mRNA levels were similar across treatments (Fig. 5G; all *P*≥0.68), while FXYD1 protein was below the limit of detection. Although FXYD5 mRNA levels remained unaltered (all *P*≥0.23), FXYD5 protein levels increased under glucose deprivation (*P*=0.02) and a similar trend (*P*=0.09) was observed under ischemic conditions (Fig. 5H,O).

### Effect of intermittent ischemic preconditioning and acute (2-hour) ischemia on HIF-1α, AMPK, and NKA in cultured human myotubes

To determine whether shorter bouts of ischemia have a different effect on NKA expression than prolonged ischemia, human myotubes were exposed to intermittent ischemia, hypoxia, or glucose deprivation for 1 hour per day for 3 days. In addition, to examine if such preconditioning affects subsequent exposure to ischemia, preconditioned myotubes were exposed to 2 hours of acute ischemia 23 hours after the last preconditioning session (see the scheme in Fig. 1B). Under ischemic conditions, HIF-1α mRNA was unchanged (Fig. 6A; *P*=0.46), while HIF-1α protein and VEGF mRNA were increased (both *P*<0.001) (Fig. 6B,C). An increase in PGK1 mRNA was marginal (*P*=0.001) (Fig. 6D). Phosphorylation of AMPK was decreased after 2 hours of acute ischemia (*P*<0.001) (Fig. 6E,F), but a marked increase in phosphorylation of ACC (*P*<0.001) indicated activation of AMPK (Fig. 6G,H). Preconditioning did not affect ischemia-induced responses of HIF-1α or AMPK (Fig. 6A-H; all *P*≥0.26).

**Fig. 6.**
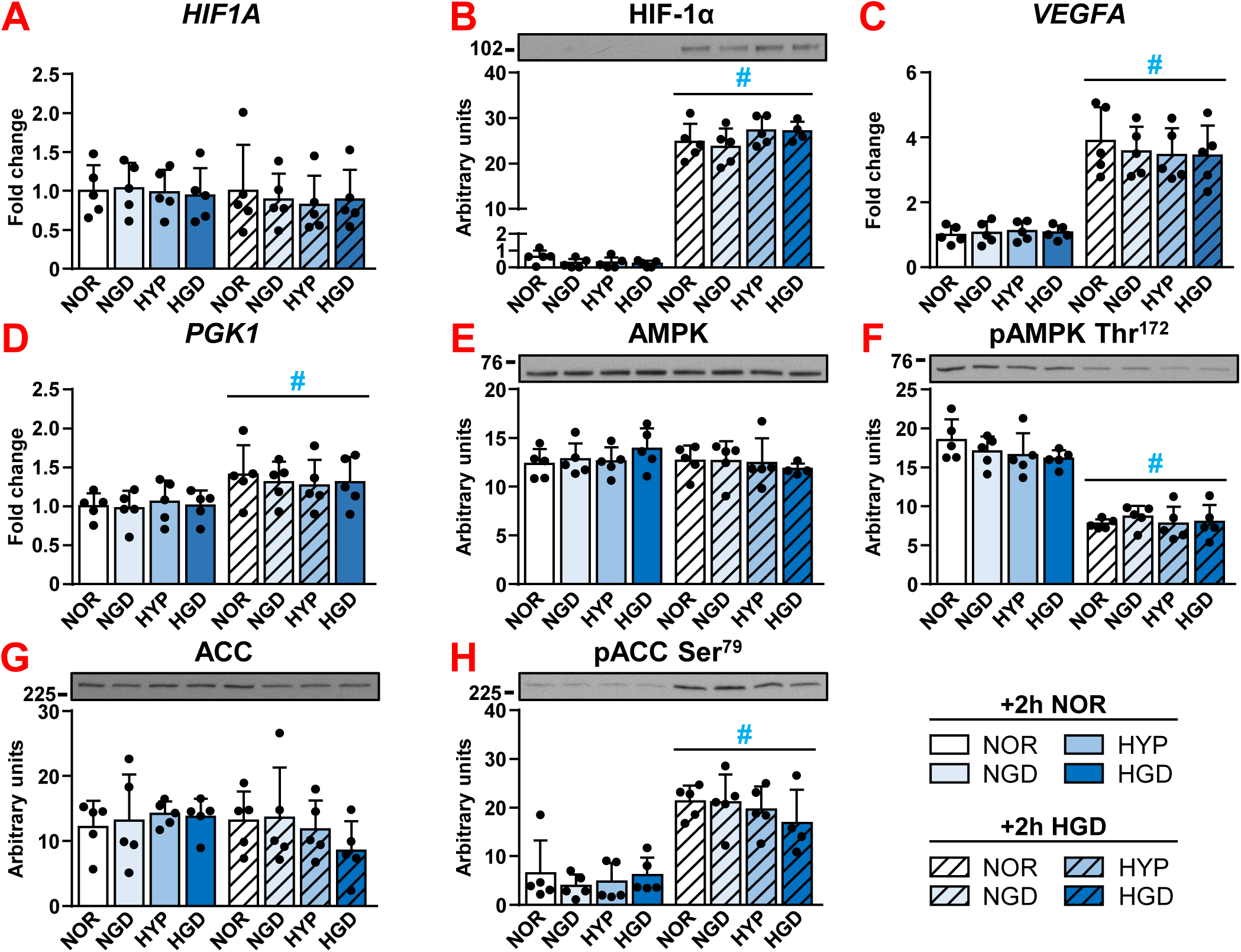
Effect of intermittent ischemic preconditioning and acute (2-hour) ischemia on HIF-1α and AMPK in cultured human myotubes. qPCR was used to estimate gene expression of (A) HIF-1α (*HIF1A*), (C) *VEGFA*, and (D) *PGK1*. 18S rRNA was used as endogenous control. Immunoblotting was used to estimate protein abundance of (B) HIF-1α, (E) AMPK, (F) pAMPK (Thr^172^), (G) ACC, and (H) pACC (Ser^79^). Results are means ± SD for n=5; \**P*<0.05 vs. NOR (+2h NOR or 2h HGD); the line with # indicates *P*<0.05 for effect of 2-hour ischemia (HGD) vs. control conditions (+2h NOR). Regular two-way ANOVA with Dunnett’s post-hoc test was used for statistical evaluation.

After 3 days of preconditioning, the mRNA expression of NKA subunits and FXYDs was similar across all conditions (Fig. 7A-H; all *P*≥0.64). Protein abundance of NKAα1 (Fig. 7I; *P*≥0.24), phosphorylated (Tyr^10^) NKAα1 (Fig. 7J; *P*≥0.30), NKAβ1 (Fig. 7M; *P*≥0.06), and FXYD5 (Fig. 7N; *P*≥0.58) was also unaltered by preconditioning. However, after 2 hours of acute ischemia, protein abundance of NKAα2 was increased in myotubes that had undergone ischemic preconditioning (*P*=0.04) (HGD, Fig. 7L). Acute ischemia also increased the overall level of Tyr10 phosphorylation (Fig. 7J; *P*=0.01), although the ratio between phosphorylated and total NKAα1 did not differ significantly (Fig. 7J; all *P*≥0.07).

**Fig. 7.**
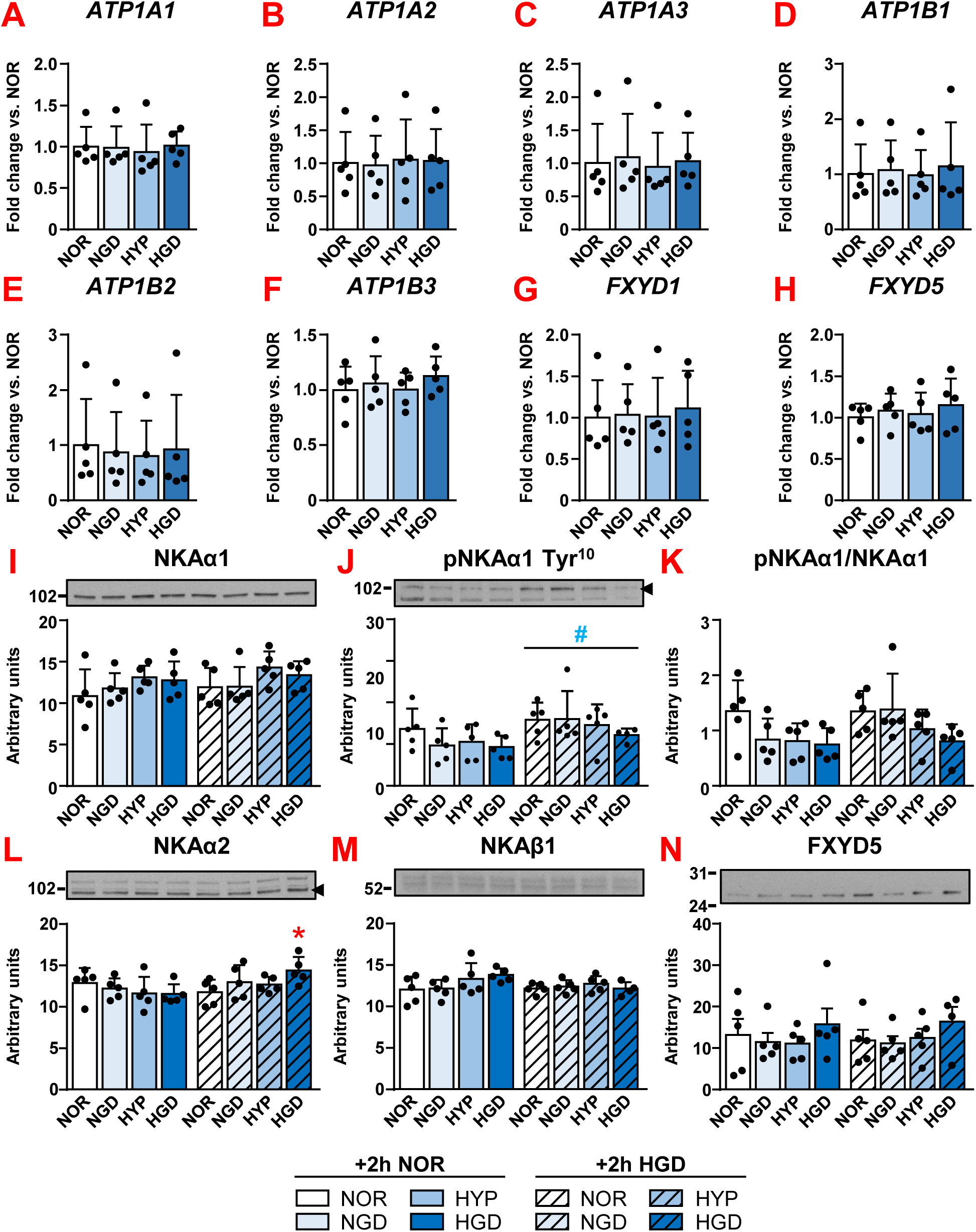
Effect of intermittent ischemic preconditioning and acute (2-hour) ischemia on the expression of NKA subunits, FXYD1, and FXYD5 in cultured human myotubes. qPCR was used to estimate gene expression of (A) NKAα1 (*ATP1A1*), (B) NKAα2 (*ATP1A2*), (C) NKAα3 (*ATP1A3*), (D) NKAβ1 (*ATP1B1*), (E) NKAβ2 (*ATP1B2*), (F) NKAβ3 (*ATP1B3*), (G) *FXYD1*, and (H) *FXYD5*. Immunoblotting was used to estimate protein abundance of (I) NKAα1, (J) pNKAα1 (Tyr^10^), (K) phosphorylation of NKAα1, relative to total protein, (L) NKAα2, (M) NKAβ1, and (N) FXYD5. Results are means ± SD for n=5; \**P*<0.05 vs. NOR (+2h NOR or 2h HGD); the line with # indicates *P*<0.05 for effect of 2-hour ischemia (HGD) vs. control conditions (+2h NOR). In case of unspecific bands arrowheads mark the specific bands that were used in densitometric analysis. For qPCR results ordinary one-way ANOVA with Dunnett’s post-hoc test was used, while regular two-way ANOVA with Dunnett’s post-hoc was used for immunoblotting results.

## DISCUSSION

Here we found that NKAα1 mRNA expression in vastus lateralis muscle was higher in subjects with ACL rupture who underwent LL-BFR exercise training than in those who underwent LL-Sham exercise training or those without intervention (Control). In addition, NKAβ3 and FXYD5 mRNA expression in vastus lateralis muscle were higher in the LL-BFR than in the LL-Sham group. However, while increases in mRNA levels support our hypothesis that LL-BFR training promotes NKA and FXYD expression in vastus lateralis of subjects with ACL rupture, without the corresponding increases in protein abundance, these transcriptional changes cannot explain functional improvements in knee extensors that we had observed in the LL-BFR group (5).

In healthy subjects, NKAα1 mRNA did not increase in vastus lateralis muscle after exercise (41). Nor were the protein levels increased after cycling bouts with BFR (12). Nevertheless, the type II fibres of vastus lateralis muscle of the leg that cycled with BFR showed higher abundance of NKAα1 than the control leg that cycled without BFR (41). In contrast to NKAα1, we found that NKAα2 was similar in the LL-BFR, LL-Sham, and Control groups, consistent with the BFR study in healthy subjects (12). Therefore, the effects of the LL-BFR exercise training on the abundance of NKAα1 and NKAα2 in skeletal muscles of subjects with knee injury appear to be similar to those observed in healthy subjects performing high-load BFR training. Interestingly, haplodeficiency of the NKAα1 subunit in mice (α1^+/-^) is associated with reduced size of soleus muscle (31), suggesting that NKAα1 may play a role in regulating muscle size. Thus, it would be interesting to further explore whether upregulation of NKAα1 mRNA is associated with the trophic effects of LL-BFR training in subjects with ACL rupture (5).

Expression of FXYD1 in vastus lateralis muscle was unaltered, which seems to exclude its involvement in adaptation of knee extensors to LL-BFR training in subjects with ACL rupture. In contrast, in healthy subjects who performed interval cycling with BFR, the abundance of FXYD1 was increased in vastus lateralis muscle (12). Running with BFR also acutely increased FXYD1 mRNA in vastus lateralis muscle of healthy subjects (41). The apparent discrepancies between our study and previous studies in healthy subjects (12, 41) remain to be explained, but might be related to differences in experimental protocols, including exercise volume and intensity, type of exercise, and timing of biopsy sampling. In addition, we analysed whole muscle samples, which may have masked fibre-type specific changes, which are known to exist (12, 41).

Expression of FXYD5 mRNA in vastus lateralis muscle was higher in the LL-BFR than in the LL-Sham group. While our study did not reveal the underlying mechanism, it is possible that ischemia caused by BFR contributed to this effect. Interestingly, previous studies showed that spinal cord injury increases FXYD5 protein abundance in vastus lateralis muscle (28), while high intensity interval training in healthy subjects decreases it in type IIa fibers from vastus lateralis muscle (64). Taken together, these studies suggested that FXYD5 content in skeletal muscle is increased by physical inactivity and decreased by physical activity. Our results in semitendinosus muscle, in which FXYD5 protein levels were reduced in the LL-BFR group, seem to better fit this pattern, but their interpretation has to be cautious because our analysis lacked statistical power due to low number of samples that were useful for immunoblotting analysis of FXYD5 (see Methods). Since LL-BFR had more pronounced functional effect on extensors than flexors (5), different molecular responses in vastus lateralis and semitendinosus muscles are not unexpected, but remain to be explained.

We used cultured myotubes to determine whether and how different components of ischemia, such as hypoxia and glucose deprivation, individually or jointly modulate NKA expression, which led us to three important observations. Firstly, prolonged ischemia and glucose deprivation had an isoform-specific suppressive effect on expression of NKA subunits. Ischemia and glucose deprivation suppressed NKAα1 mRNA and protein levels, but had no effect on NKAα2, which indicates fundamental differences in regulation of the two major muscle NKAα isoforms. The abundance of NKAα1 is increased in cultured human myotubes under high glucose conditions (48) and in choroid plexus of hyperglycaemic diabetic rats (65), while NKAα1^+/-^ mice seem to have reduced capacity for glucose oxidation in the hearts (66). Taken together, these studies suggest a close link between NKAα1 and glucose metabolism, but they do explain the upregulation of NKAα1 and NKAβ3 mRNA in the vastus lateralis in the LL-BFR group.

Secondly, the abundance of FXYD5 was increased by glucose deprivation, which indicates that the regulation of FXYD5 is linked to energy metabolism. We suspect that increased FXYD5 abundance with a concurrent decrease in NKAα1 and NKAβ1 abundance would results in altered composition of NKA α/β-heterodimers and NKA/FXYD complexes, thus affecting their kinetic properties (7, 8, 19, 21, 25). For instance, FXYD5 associates with NKA and increases its *V_max_* (58, 67), suggesting that the upregulation of FXYD5 might be an adaptive response to preserve transmembrane transport of Na^+^ and K^+^ despite downregulation of NKA. Interestingly, FXYD5 has been associated with a negative regulation of NKAβ1 expression in cultured 4T1 cancer cells (68). The downregulation of NKAβ1 in myotubes under glucose deprived conditions could therefore be a consequence of the increased abundance of FXYD5. Although any extrapolations from cell culture experiments to the *in vivo* situation have limited validity, it has to be emphasized that upregulation of FXYD5 protein was the only *in vitro* result that, at least to some extent, mimicked what we found in vastus lateralis muscle (upregulation of FXYD5 mRNA in the LL-BFR group). It would therefore be interesting to further explore the role of glucose and/or ischemia in regulation of FXYD5 in skeletal muscle.

Thirdly, hypoxia alone had no effect on NKA or FXYD expression in myotubes. Conversely, glucose deprivation almost completely replicated the effect of ischemia on NKA expression. Changes in NKA expression in ischemic myotubes therefore seem to be almost completely driven by glucose deprivation. Previous studies investigating the role of hypoxia on NKA in skeletal muscle produced inconsistent results. Climbing in normobaric or hypobaric hypoxia reduced the concentration of NKA in vastus lateralis muscle of healthy subjects (69, 70). However, during live high, train low hypoxia intervention, hypoxia did not alter the NKA content in vastus lateralis muscle (71). Running under systemic hypoxia also did not acutely alter expression of NKA subunits (except NKAβ1) or FXYD1 in vastus lateralis of healthy subjects (41). In addition, while hypoxia had no effect on FXYD5 in cultured myotubes, it downregulated FXYD5 in murine macrophages (72) and upregulated FXYD5 in the carotid body (73). Clearly, effect of hypoxia on NKA and FXYDs is complex and depends on several factors, such as cell type and experimental protocol.

Phosphorylation is a major mechanism by which the subcellular distribution and activity of NKA in skeletal muscle is regulated (21). Here we determined that LL-BFR training reduced the ratio between the phosphorylated (Tyr^10^) and total NKAα1 in semitendinosus muscle, which may suggest that NKAα1 Tyr^10^ is responsive to changes in the muscle metabolic status. This notion is supported by changes in the level of phosphorylated (Tyr^10^) NKAα1 in cultured myotubes induced by prolonged and acute 2-hour ischemia. In the kidney, phosphorylation of Tyr^10^ is increased by insulin, which stimulates NKA activity (74, 75). Conversely, the dephosphorylation of Tyr^10^ was linked to increased NKA activity in skeletal muscle (76). We had previously shown that activation of AMPK in kidney cells suppressed the phosphorylation of Tyr^10^ (46), but in the present study, AMPK activation does not appear to correlate with the phosphorylation status of Tyr^10^ in either vastus lateralis or semitendinosus muscle. Similarly, although AMPK activation was associated with a decrease in the level of phosphorylated Tyr^10^ in cultured myotubes exposed to prolonged ischemia, the concomitant decrease in the total amount of NKAα1 makes the mechanistic interpretation of these alterations challenging.

### Limitations

One of the strengths of the study is that we examined the effects of LL-BFR training on the expression of NKA subunits and FXYDs in skeletal muscles of subjects with an ACL injury, which had not been done before. We also examined both the extensor (vastus lateralis) and the flexor (semitendinosus) muscles, which is rarely done because semitendinosus muscle is less accessible (77) for standard needle biopsy than vastus lateralis muscle. Furthermore, we used cultured myotubes to provide new insights into molecular mechanisms that link ischemia to regulation of NKA expression and phosphorylation. However, our study also has several weaknesses. Firstly, in order to minimize unnecessary interventions unrelated to the treatment of the ACL injury, muscle biopsies were obtained exclusively from the injured limb during ACL reconstruction surgery. The collection of samples at the beginning of the study (i.e. before the start of training) and/or from the contralateral (uninjured) leg would have increased the validity of our study. Multiple sampling would also optimize our ability to detect changes in mRNA and protein expression after exercise, which typically occur with different timeframes and dynamics (78) and may have been missed only because we collected samples at a single time point.

Secondly, vastus lateralis biopsies were small, which limited the amount of protein that was available for immunoblotting, while the tendinous part of semitendinosus muscle apparently predominated in some samples, so that they had to be excluded from the analysis due to the low content of myosin. Clearly, relatively low number of samples combined with significant biological variability and relatively low sensitivity of the Western blot limited our ability to detect small to moderate alterations in the abundance of NKA in skeletal muscle (79).

Thirdly, cultures of myotubes were prepared from semitendinosus muscles of subjects with ACL rupture. Such approach has the advantage of using surgical waste obtained during ACL surgery and was previously used by us (53, 56, 57, 80–82) and others (83–85) not only to study skeletal muscle cells, but also non-muscle cells (86–89) as well as skeletal muscle transcriptome (90) and proteome (86, 87, 91). A potential disadvantage of our approach is that rupture of ACL and subsequent knee dysfunction might have affected satellite cells (92) in semitendinosus muscle and consequently the experimental results obtained in myotubes derived from these cells. A systematic comparison of myotubes from healthy and injured subjects would clarify this question, but would be difficult to perform for ethical reasons. Without such comparison we cannot exclude the possibility that our *in vitro* results were affected by the ACL rupture. However, even if we assume that ACL rupture altered the properties of the myotubes we used, it could be argued that our approach would be valid for studying effects of ischemia on NKA expression in skeletal muscle affected by the knee injury.

Finally, our study did not include a group that would be subjected to BFR without exercise, which would enable us to study effects of ischemia on NKA and FXYD expression *in vivo*. Using cultured myotubes to dissect mechanism linking ischemia and NKA expression has obvious disadvantages, not least because conditions during LL-BFR training cannot be directly mimicked *in vitro*. For instance, non-innervated primary human myotubes, which we used in these experiments, are quiescent and do not contract spontaneously if not electrically stimulated (93–95), so we could not explore the effect of ischemia and contractions simultaneously. Another disadvantage in the context of NKA research is that expression of the NKAα1 subunit is relatively more prominent in cultured myotubes than in skeletal muscle tissue (21, 53, 56). Nevertheless, cultured human myotubes are a useful and valid model to study different aspects of skeletal muscle function (96, 97), especially molecular mechanisms and treatments that are difficult or impossible to study experimentally in humans.

### Conclusions and perspective

Studies in healthy subjects indicated that isoform-specific alterations in the expression of NKA subunits and/or FXYDs may be involved in muscle adaptations in response to BFR training (12, 13, 41). However, while ACL rupture was shown to impair NKA in vastus lateralis muscle (2), data on the effect of LL-BFR training on NKA expression in subjects with ACL injury, who may particularly benefit from LL-BFR exercise training, were lacking. Our study now fills this gap by providing evidence that 3-weeks of LL-BFR exercise training is associated with increased NKAα1, NKAβ3, and FXYD5 mRNA expression in vastus lateralis muscle in subjects with ACL rupture. Without changes in NKA protein abundance, these results cannot explain improved function of knee extensors in these subjects (5). Nevertheless, they provide a basis for furtherer exploration of the role of NKA in muscle adaptations to LL-BFR training after ACL rupture. To further dissect the role of BFR, future studies should also compare effects of LL-BFR exercise and BFR without exercise.

We also showed that prolonged ischemia, but not intermittent ischemia, reduced the mRNA levels of NKAα1, NKAβ1, and NKAβ3 and protein levels of NKAα1 and NKAβ1 in cultured myotubes. These experiments do not explain upregulation of NKAα1 and NKAβ3 mRNA due to LL-BFR training, but show that ischemia exerts isoform-specific effects on NKA expression in myotubes mainly via glucose deprivation rather than hypoxia. Conversely, our results suggest that glucose deprivation may promote upregulation of FXYD5. Future studies should investigate whether concurrent ischemia and myotube contractions produce a different effect on NKA or FXYD expression.

## DATA AVAILABILITY STATEMENT

Raw data for immunoblots and PCR that support the findings of this study are available in the supplemental material of this article.

## SUPPLEMENTAL MATERIALS

Supplemental materials include raw data of immunoblotting (Supplement 1) and real-time PCR (Supplement 2).

## ACKNOWLEDGEMENTS

Technical assistance of Ms Ksenja Babič (Faculty of Medicine, University of Ljubljana) is gratefully acknowledged.

## USE OF ARTIFICIAL INTELLIGENCE

A completed, fully written version of the manuscript, which had been prepared without the use of AI, was subjected to assessment by the Nature research assistant with the purpose of evaluating English grammar and clarity of the text. Suggestions by the assistant were taken into consideration, but all writing was done by the authors.

## AUTHOR CONTRIBUTIONS

**V.J.:** analysed data, performed experiments, interpreted results of experiments, prepared figures, drafted manuscript, edited and revised manuscript, approved final version; **K.M.**: analysed data, performed experiments, approved final version; **A.K:** conceived and designed research, interpreted results of experiments, edited and revised manuscript, approved final version. **A.V.**: analysed data, performed experiments, approved final version, **T.T.Ž.:** performed experiments, edited and revised manuscript, approved final version; **K.S.:** performed experiments, edited and revised manuscript, approved final version; **M.P.:** interpreted results of experiments, edited and revised manuscript, approved final version; **T.M.:** interpreted results of experiments, edited and revised manuscript, approved final version, **M.D.:** conceived and designed research, interpreted results of experiments, edited and revised manuscript, approved final version; **A.V.C:** conceived and designed research, interpreted results of experiments, edited and revised manuscript, approved final version; **S.P.:** conceived and designed research, analysed data, prepared figures, interpreted results of experiments, drafted manuscript, edited and revised manuscript, approved final version.

