## Supplement 1 for "Low-load blood flow restriction training and ischemia modulate expression of Na^+^,K^+^-ATPase and FXYDs in human skeletal muscle"

### RAW DATA OF IMMUNOBLOTTING Figs. 2-7

#### Effect of low-load exercise with blood flow restriction and in vitro ischaemia on $Na^+$,$K^+$-ATPase expression in skeletal muscle and cultured myotubes of subjects with ACL rupture


Vid Jan[1], Katarina Miš[1], Alan Kacin[2], Anja Vidović[1], Tina Tomc Žargi[2], Klemen Stražar[3], Matej Podbregar[1,4], Tomaž Marš[1], Matej Drobnič[3,5], Alexander V. Chibalin[6,7], Sergej Pirkmajer[1,*]

[1]University of Ljubljana, Faculty of Medicine, Institute of Pathophysiology, Ljubljana, Slovenia

[2]University of Ljubljana, Faculty of Health Sciences, Department of Physiotherapy, Ljubljana, Slovenia

[3]University Medical Centre Ljubljana, Department of Orthopedics, Ljubljana, Slovenia

[4]General and Teaching Hospital Celje, Celje, Slovenia

[5]Department of Orthopedics, Faculty of Medicine, University of Ljubljana, Ljubljana, Slovenia

[6]Karolinska Institutet, Department of Molecular Medicine and Surgery, Integrative Physiology, Stockholm, Sweden

[7]National Research Tomsk State University, Tomsk, Russia.

### Vastus lateralis samples

#### Figures 2+3

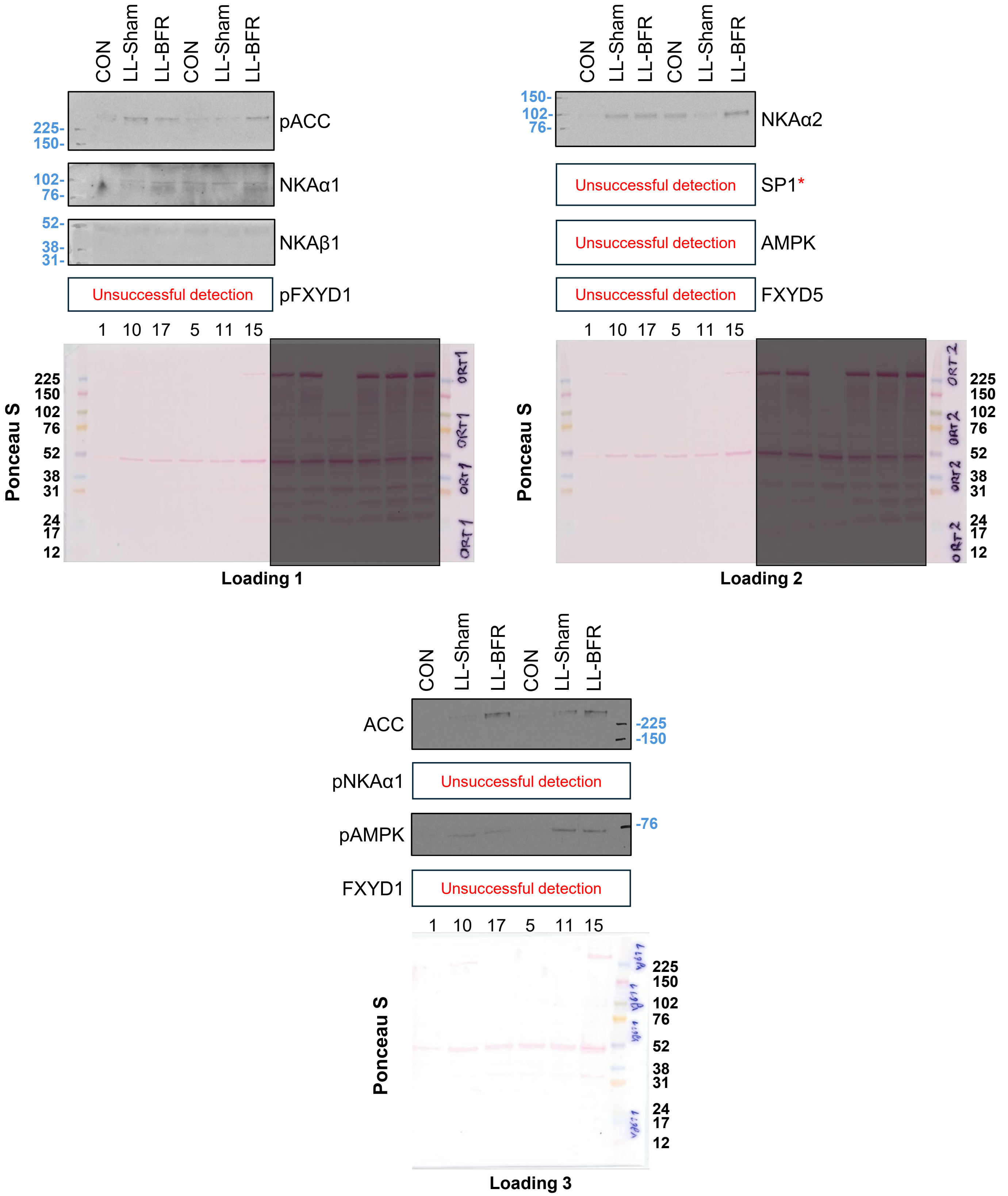


1, 10, 17, 5, 11, and 15 are sample tags for different subjects.
* labels blots that were performed on stripped membranes.

### Vastus lateralis samples

### Figures 2+3

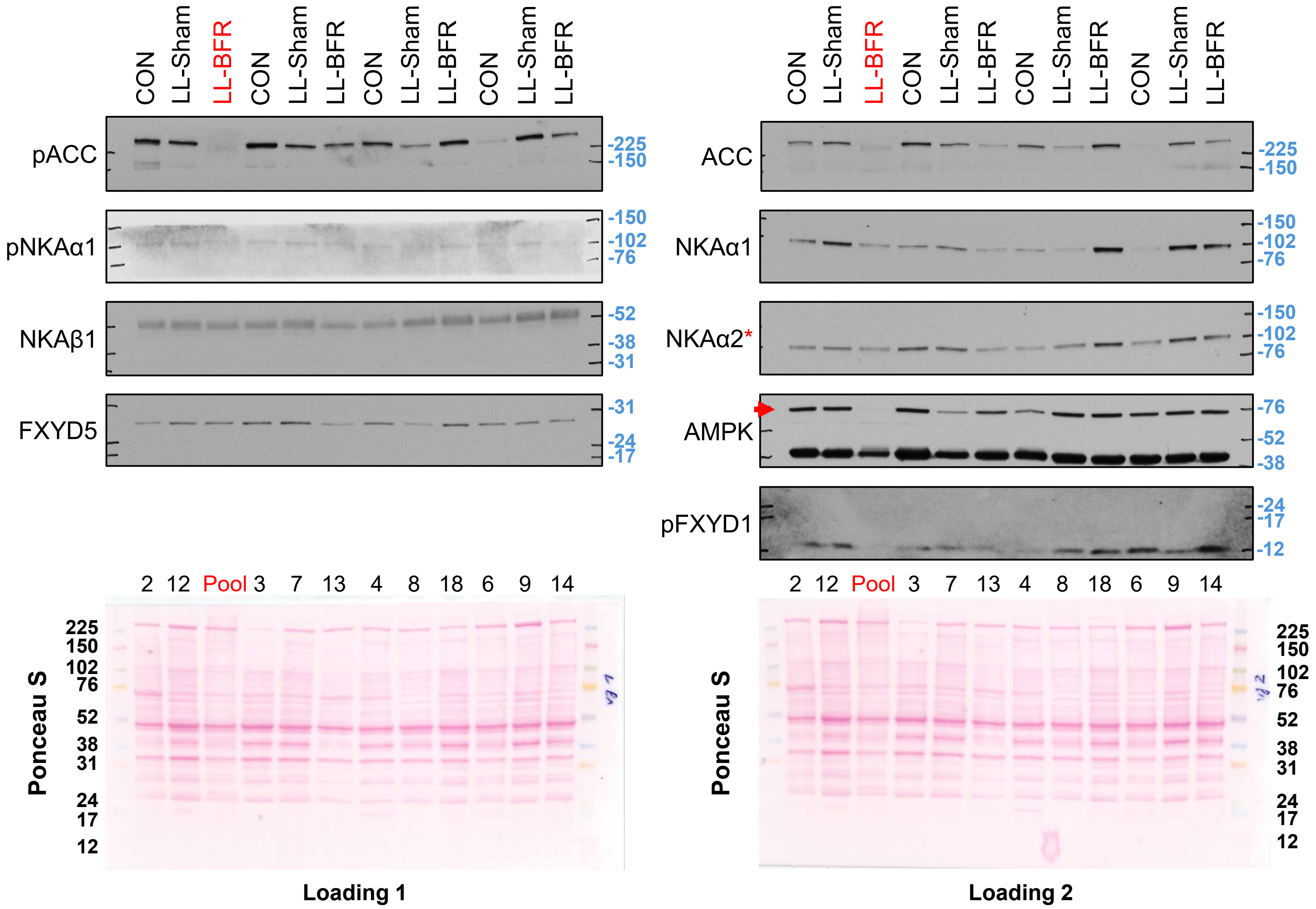


Pool sample was a pooled replacement for a missing sample and was thus excluded from statistical analysis.

2, 12, 3, 7, 13, 4, 8, 18, 6, 9, and 14 are sample tags.

* labels blots that were performed on stripped membranes. . In case of unspecific bands red arrowhead marks the specific bands that were used in densitometric analysis

### Vastus lateralis samples

### Figures 2+3

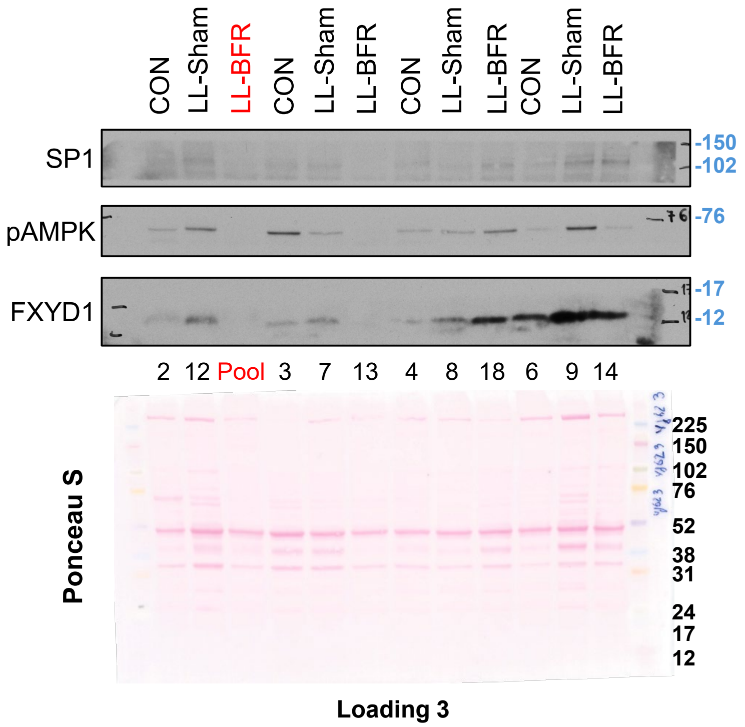


Pool sample was a pooled replacement for a missing sample and was thus excluded from statistical analysis.

2, 12, 3, 7, 13, 4, 8, 18, 6, 9, and 14 are sample tags..

### Semitendinosus samples

### Figures 2+3

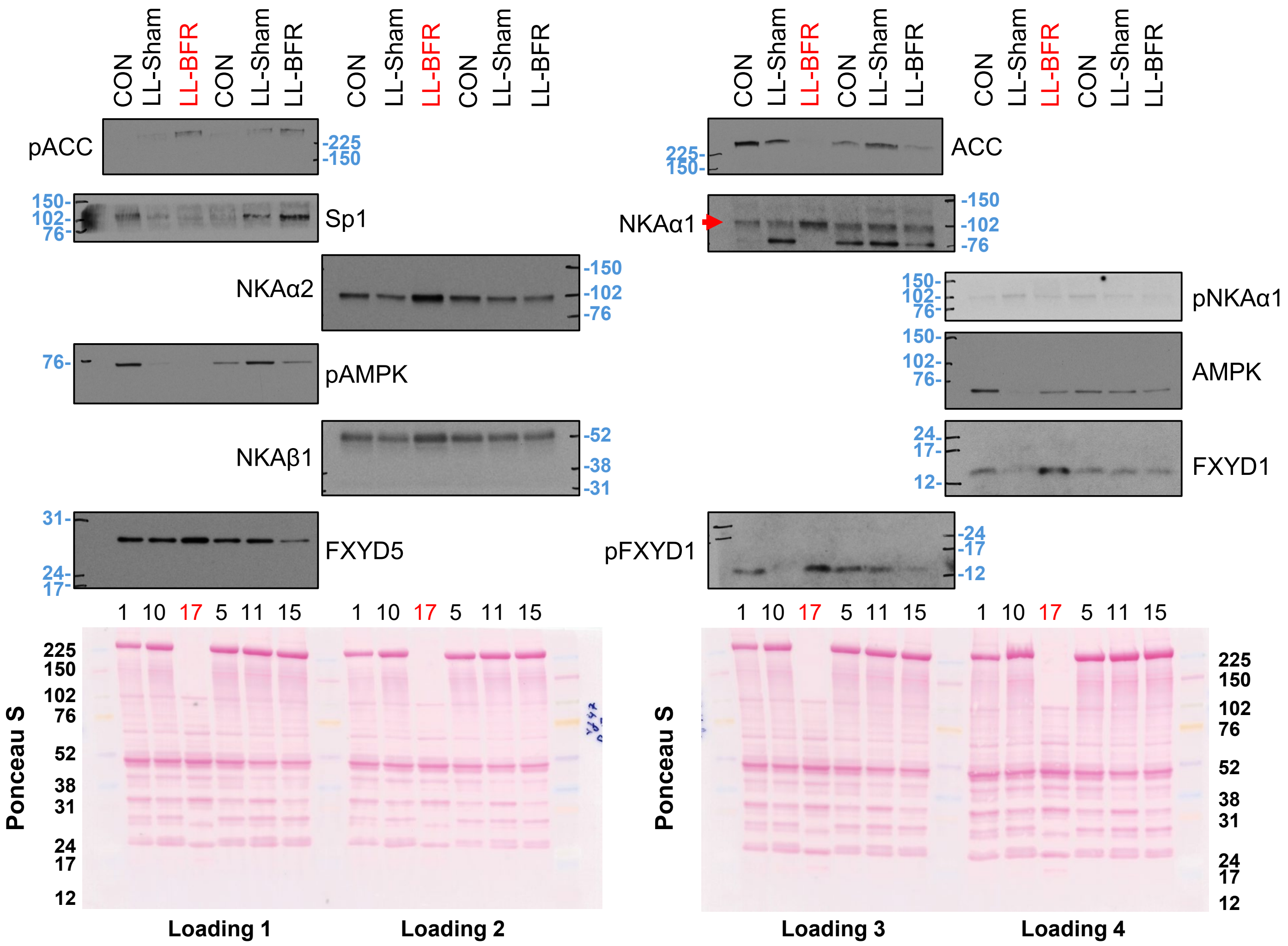


Samples from subject 17 were excluded from statistical analysis because of strange Ponceau staining.

1, 10, 17, 5, 11, 15, 2 , 12 , 16, 3, 7, 4, 8, 18, 6, 9, and 14 are sample tags.

. In case of unspecific bands red arrowhead marks the specific bands that were used in densitometric analysis

### Semitendinosus samples

### Figures 2+3

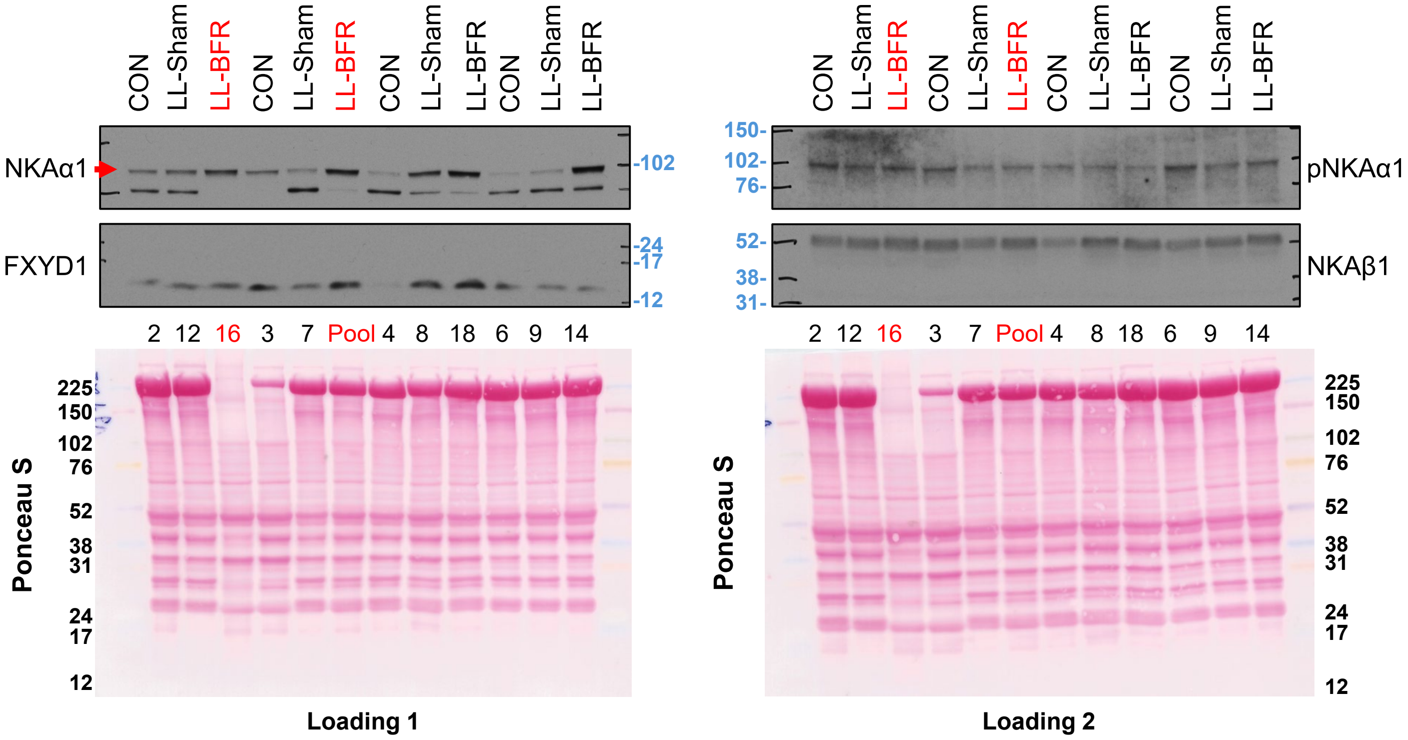


Samples from subject 16 were excluded from statistical analysis because of strange Ponceau staining. Pool sample was a pooled replacement for a missing sample and was thus excluded from statistical analysis.

1, 10, 17, 5, 11, 15, 2 , 12 , 16, 3, 7, 4, 8, 18, 6, 9, and 14 are sample tags.

. In case of unspecific bands red arrowhead marks the specific bands that were used in densitometric analysis

### Semitendinosus samples

**Figures 2+3**

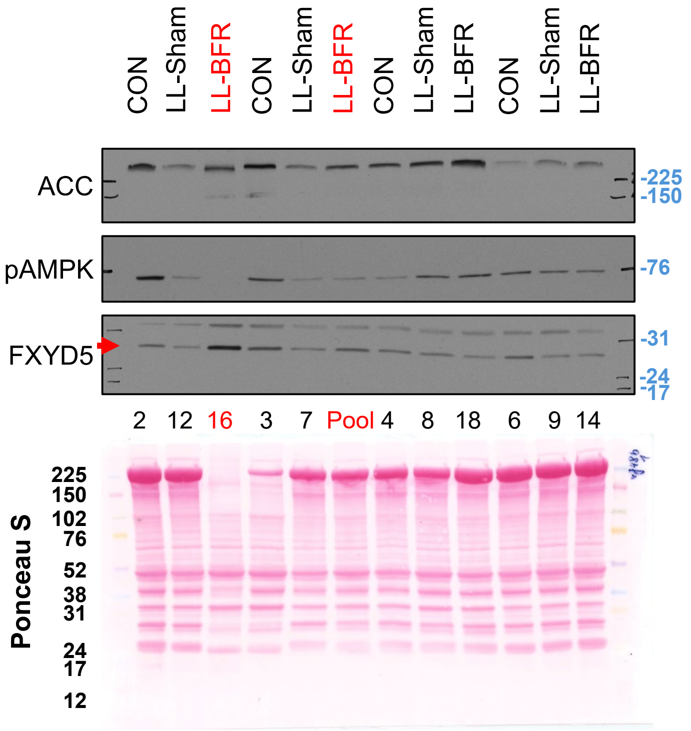


**Loading 3**

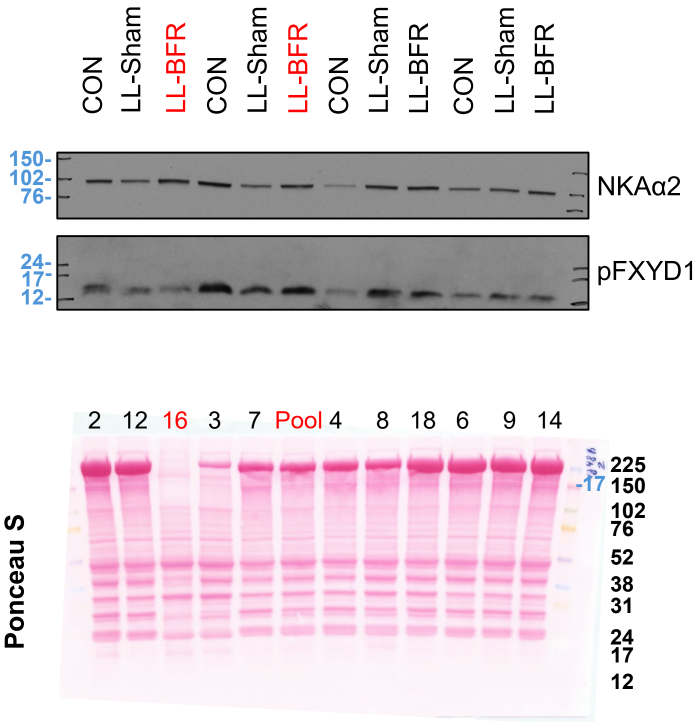


**Loading 4**

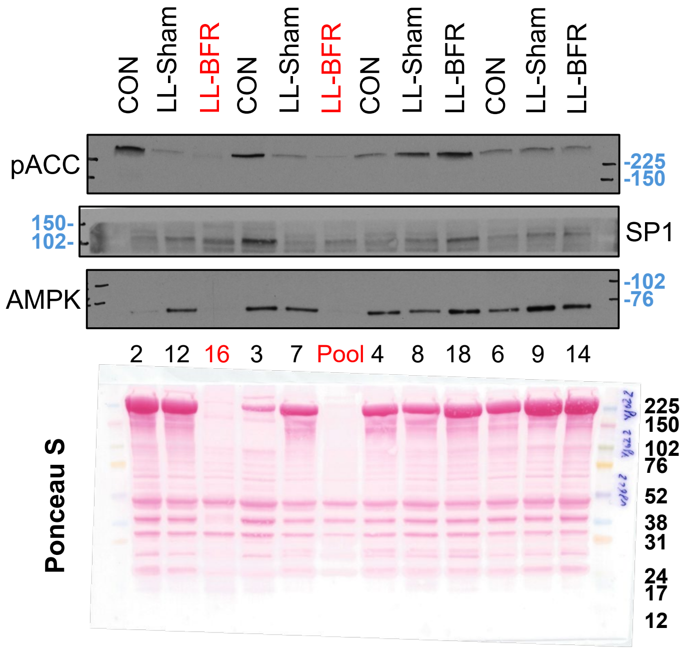


**Loading 5**

Sample 16 was excluded from statistical analysis because of strange Ponceau staining. Pool sample was a replacement for a missing sample and was thus excluded from statistical analysis.

2, 12, 16, 3, 7, 4, 8, 18, 6, 9, and 14 are sample tags.

### Figures 4+5

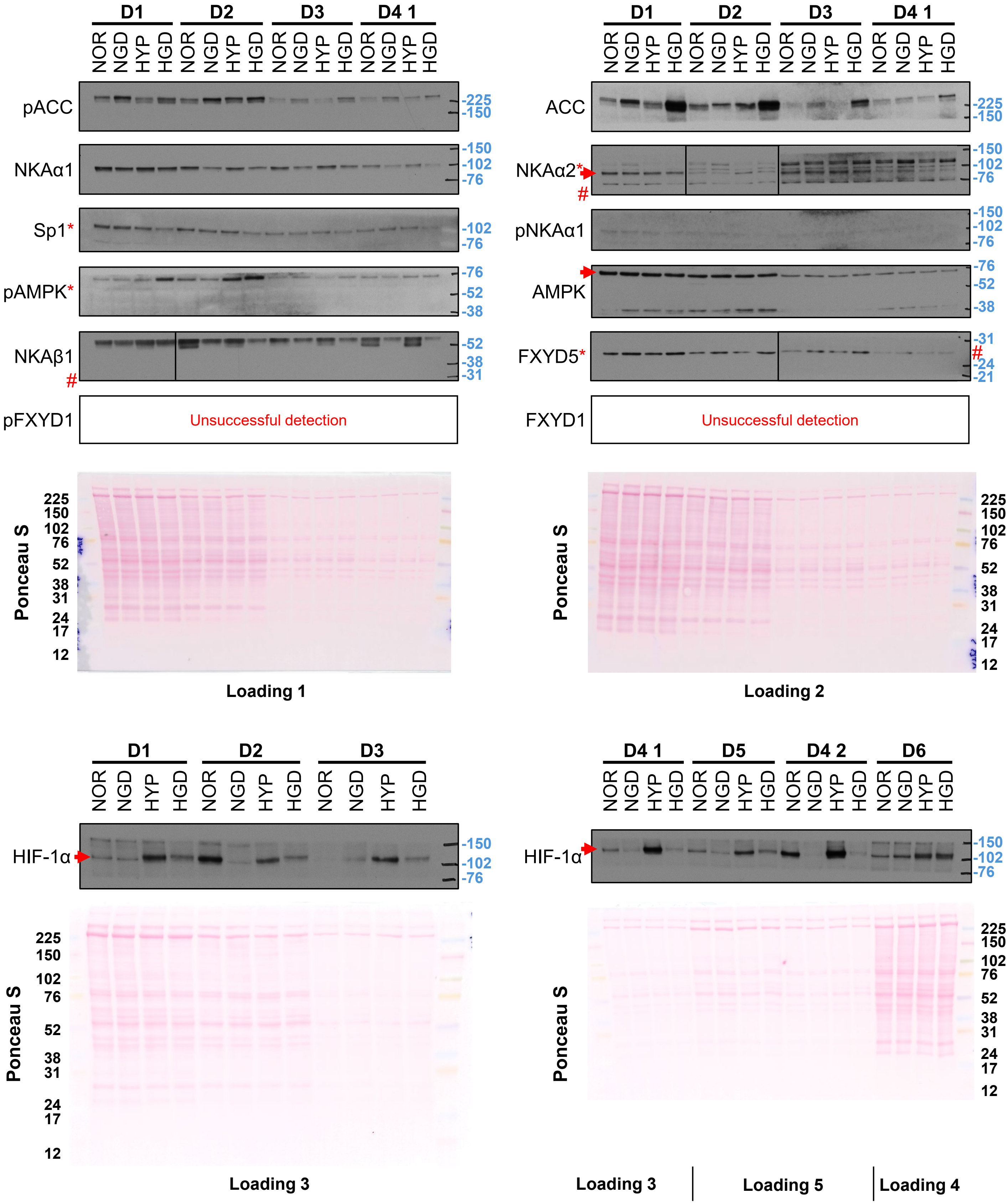


D1-6 label samples from 6 different donors, D4 1 and D4 2 were cells from the same donor, used in 2 separate experiments, so we averaged blot values

*labels blots that were performed on stripped membranes, # labels detections where due to high difference in band intensities between donors, 2 or 3 different time exposures are shown next to one another, separated by black frame.

In case of unspecific bands red arrowhead marks the specific bands that were used in densitometric analysis

### Figures 4+5

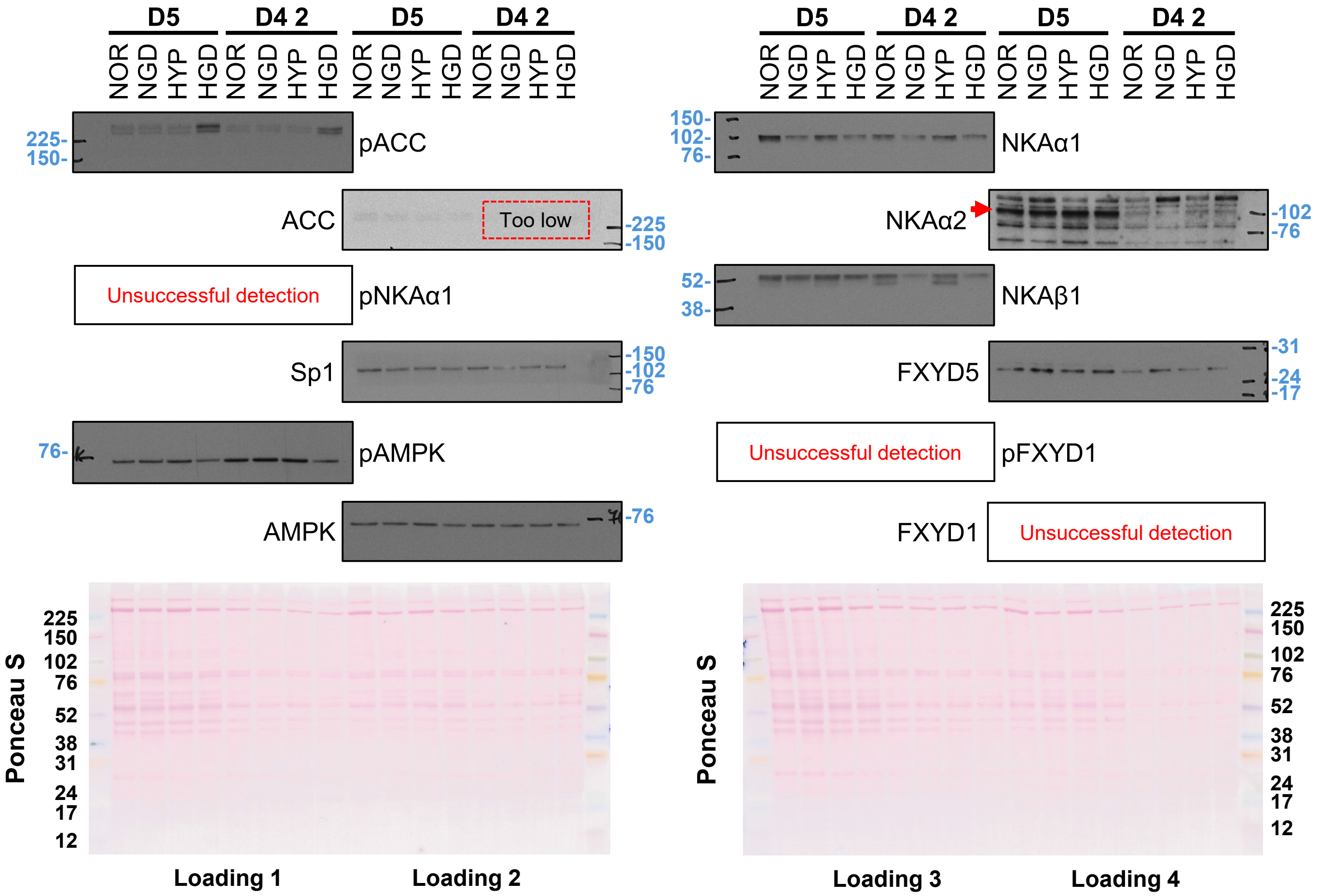


D4-6 label samples from 3 different donors, D4 1 and D4 2 were cells from the same donor, used in 2 separate experiments, so we averaged blot values

In case of unspecific bands red arrowhead marks the specific bands that were used in densitometric analysis

### Figures 4+5

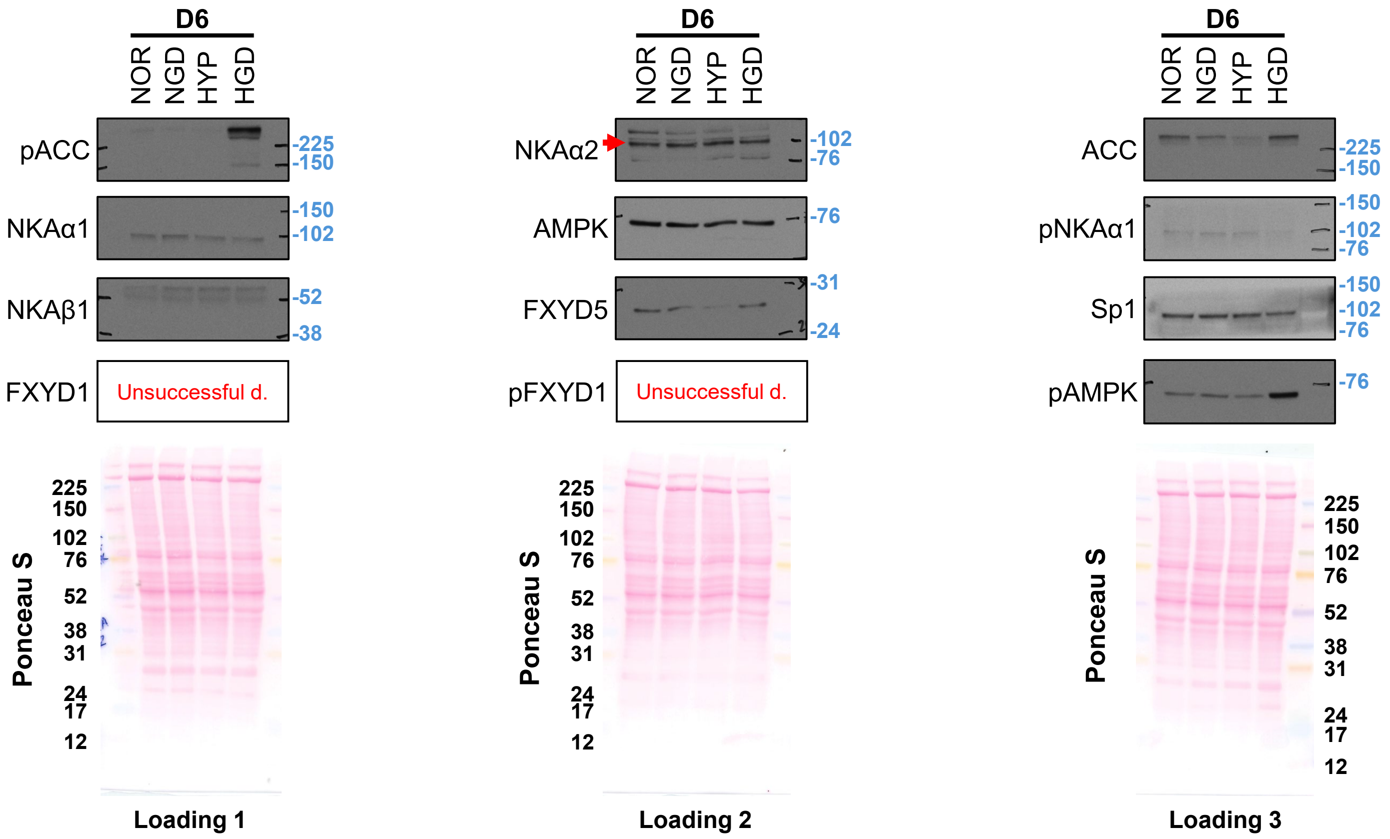


D4-6 label samples from 3 different donors, D4 1 and D4 2 were cells from the same donor, used in 2 separate experiments, so we averaged blot values

In case of unspecific bands red arrowhead marks the specific bands that were used in densitometric analysis

### Figures 6+7

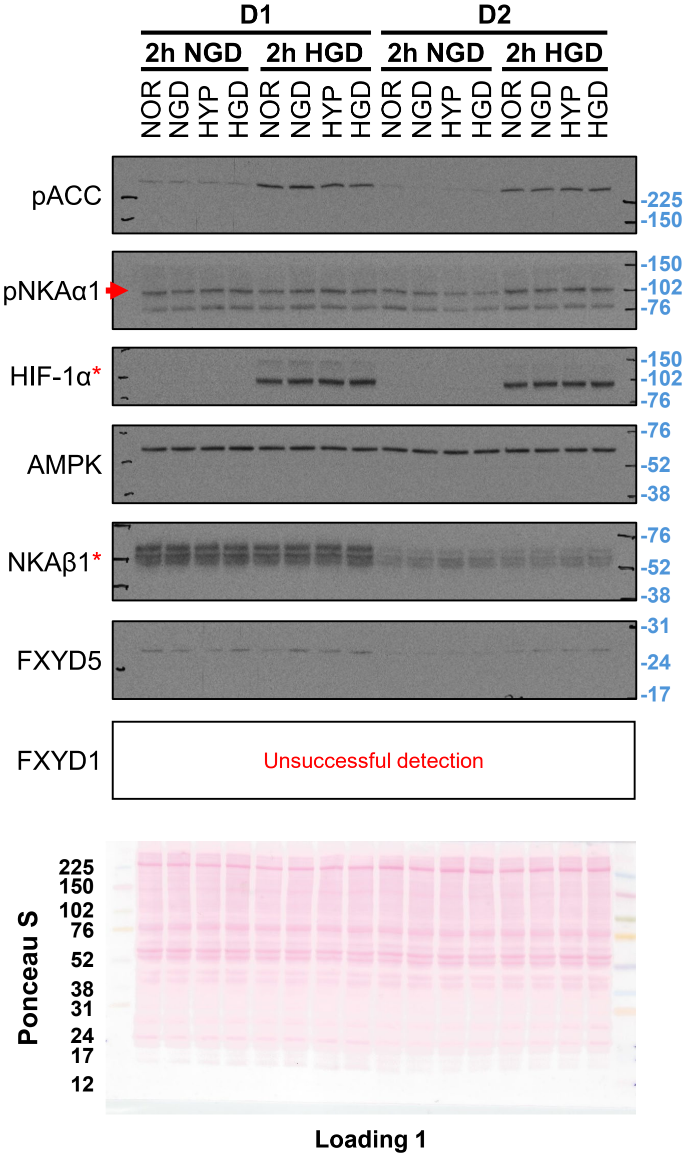


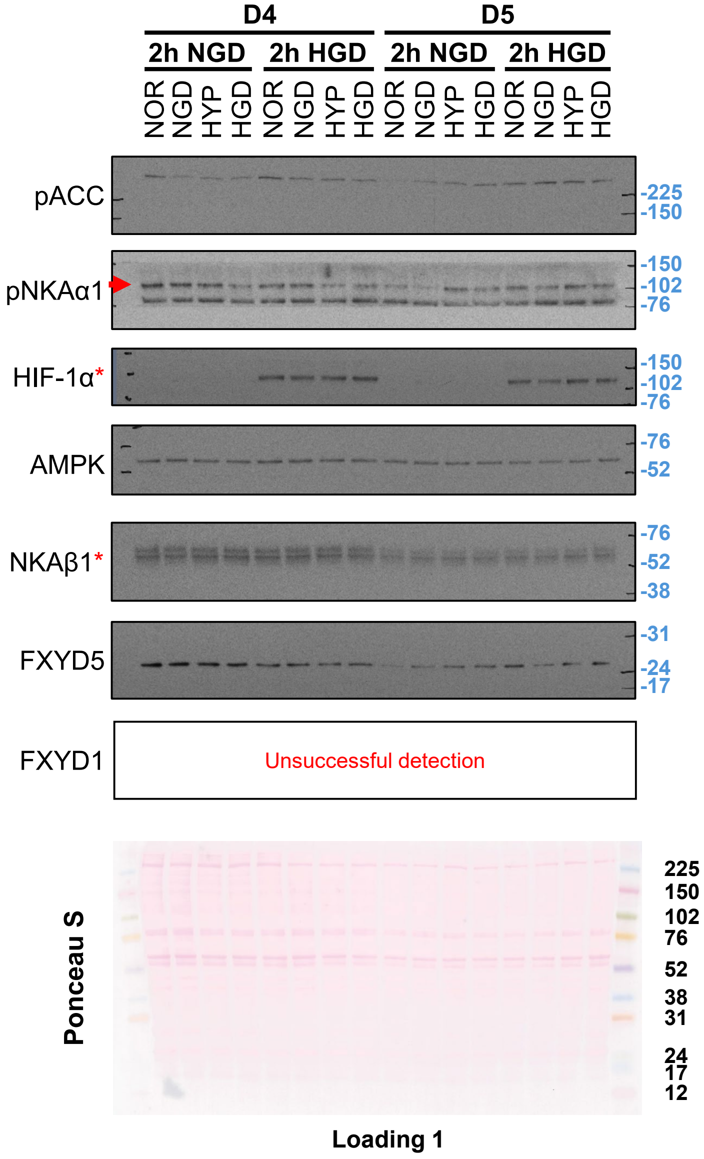


D1-5 label samples from 5 different donors

*labels blots that were performed on stripped membranesIn case of unspecific bands red arrowhead marks the specific bands that were used in densitometric analysis

### Figures 6+7

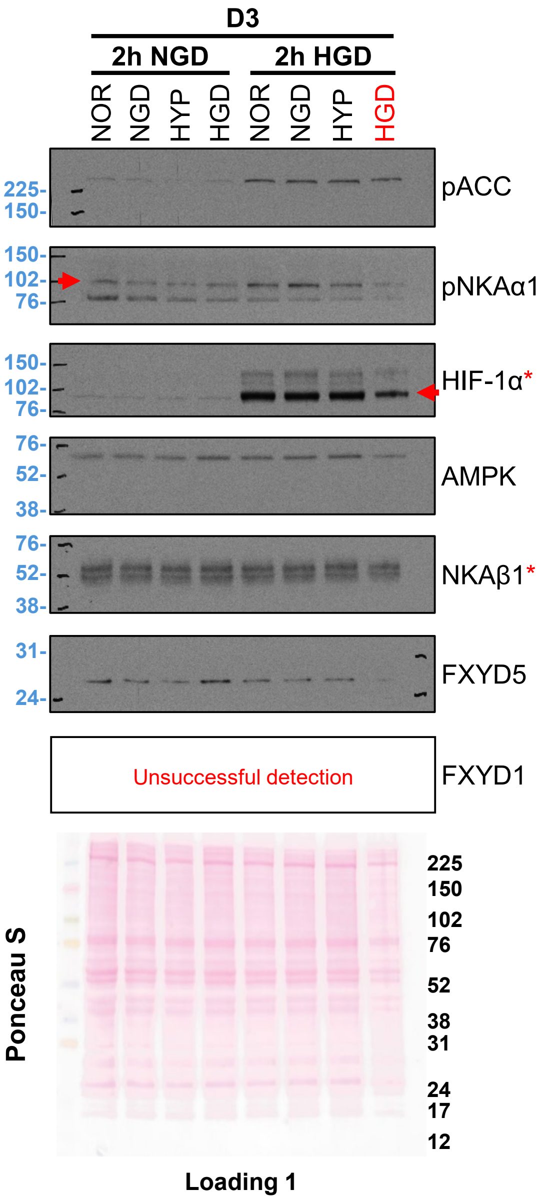


D1-5 label samples from 5 different donors
*labels blots that were performed on stripped membranesIn case of unspecific bands red arrowhead marks the specific bands that were used in densitometric analysis

### Figures 6+7

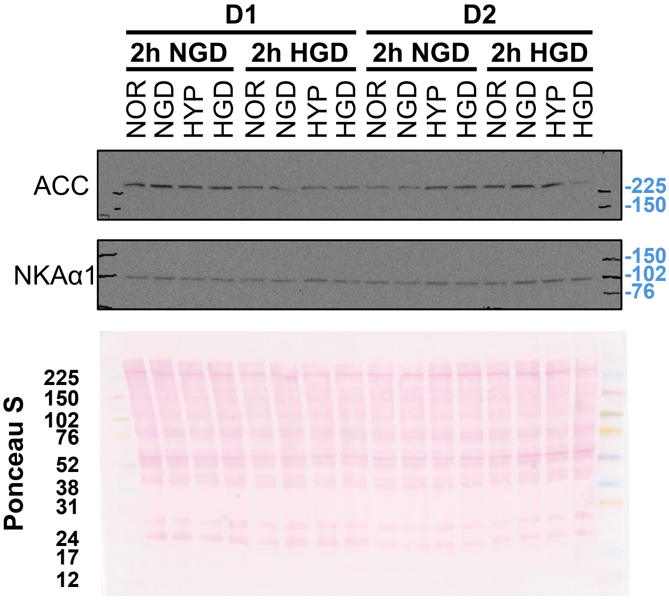


Loading 2

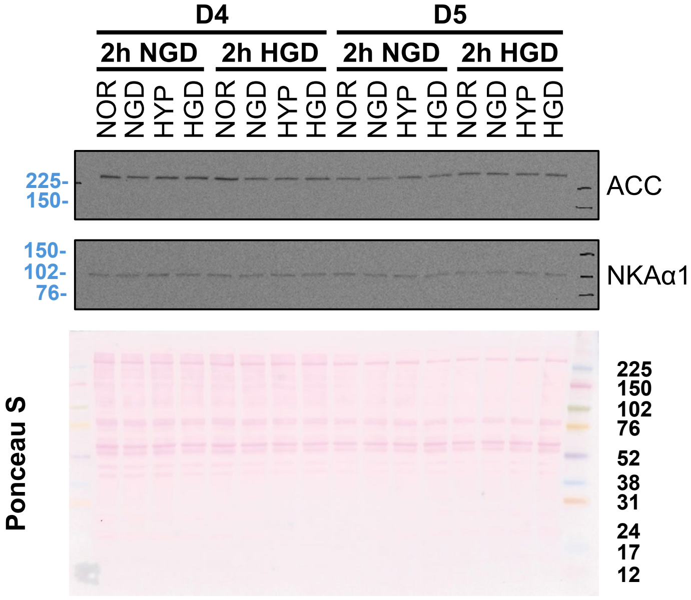


Loading 2

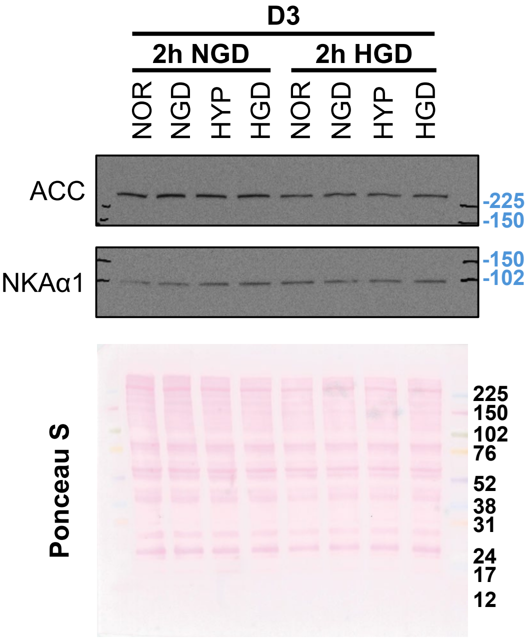


Loading 2

D1-5 label samples from 5 different donors

### Figures 6+7

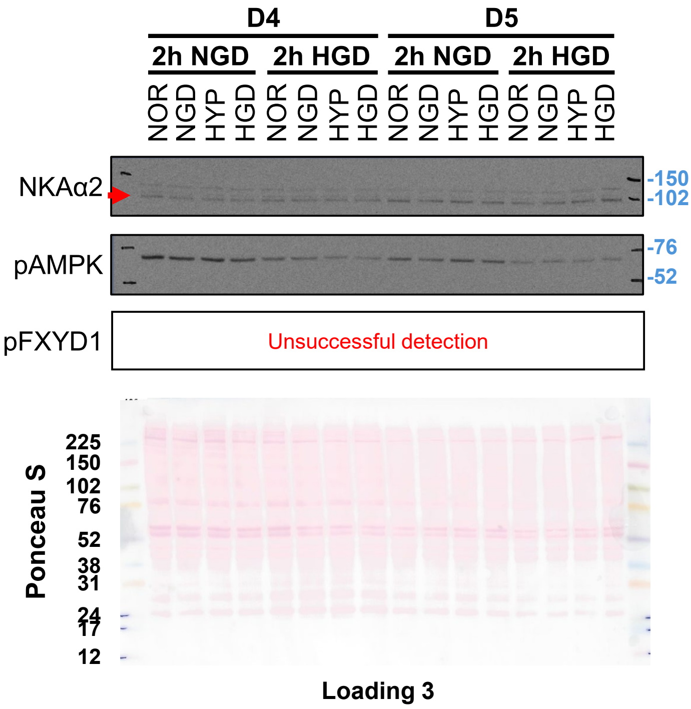


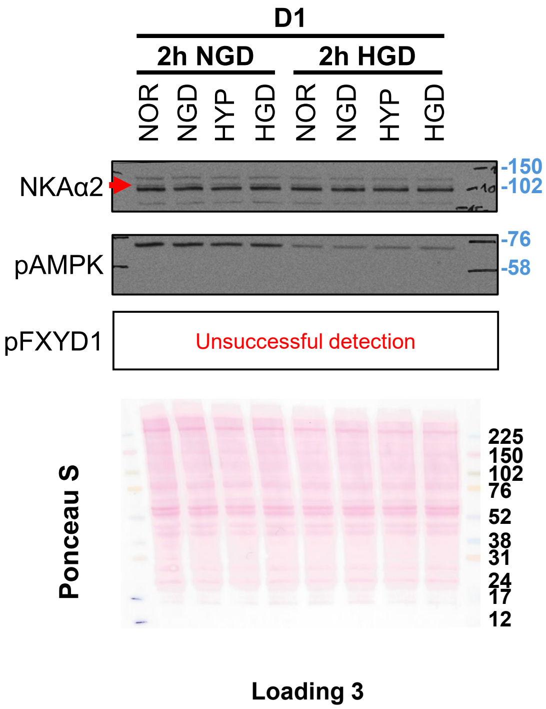


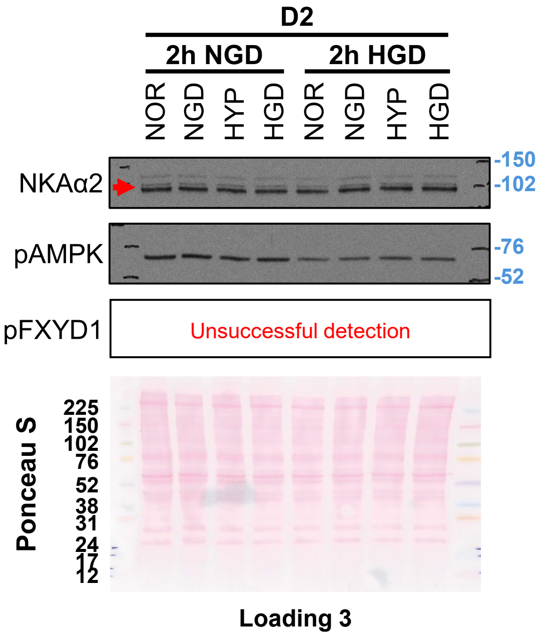


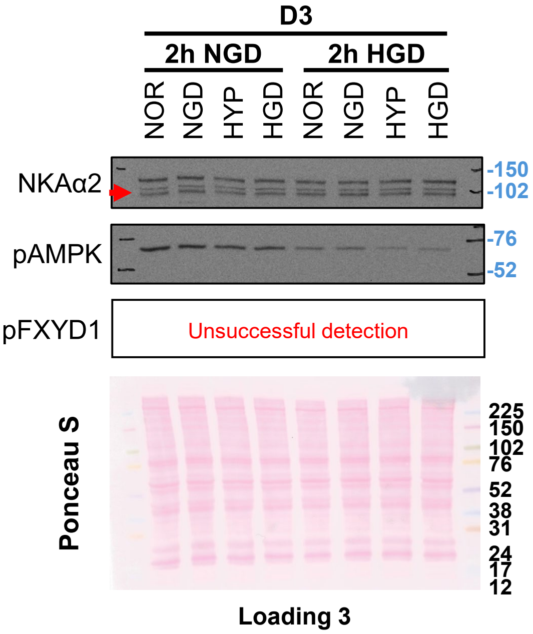


D1-5 label samples from 5 different donors

In case of unspecific bands red arrowhead marks the specific bands that were used in densitometric analysis
